# Predictive Grid Cell Coding Does Not Require Explicit Planning

**DOI:** 10.64898/2026.10.09.757963

**Authors:** Aarav Sinha, Oluwajomiloju C. Fasehun, William T. Redman

**Affiliations:** Obra D. Tompkins High School, USA; Department of Neuroscience, Johns Hopkins University, USA; Department of Electrical and Computer Engineering, Johns Hopkins University, USA; Data Science and AI Institute, Johns Hopkins University, USA

## Abstract

Predictive grid cells, a recently discovered functional class of neurons in medial entorhinal cortex (MEC), exhibit grid-like tuning, not with respect to where an animal currently is, but where it will be. Such coding, which is disjoint from standard grid cell coding, appears consistent with the hypothesis that MEC supports predictive computations for goal-directed navigation. However, given that existing predictive coding theories have been used to understand standard grid cells, it remains unclear what the functional significance of predictive grid cells is. Furthermore, the extent to which predictive grid cells are generally present in MEC is unknown. To address these open questions, we analyze data from rodent electrophysiology recordings and recurrent neural networks (RNNs). In an independent, previously recorded MEC dataset, we find evidence of predictive grid cell coding when the animal is awake and exploring an open field environment, suggesting that they are a general feature of rodent MEC. In RNNs trained to path-integrate, which have previously been shown to develop units with grid-like representations, we also find units with predictive grid-like representations, even though the RNNs are not trained with a predictive objective. We validate the predictive grid tuning of these units by finding that they exhibit significant future path-dependence, even when controlling for the same allocentric spatial position. In addition, we find that band-like units, which have been found to play an important role in maintaining path-integration in the RNNs, exhibit greater predictive grid tuning than expected by chance. These results lead us to hypothesize that predictive grid cells are a property of the underlying grid cell circuit. We test this by looking for signatures of predictive grid coding in the MEC electrophysiological data collected during sleep, finding evidence in one recording. Collectively, our results challenge the idea that predictive grid cells must necessarily be used for explicit prediction and planning, and instead may be involved in shaping the population level activity of grid cells.

## Introduction

Medial entorhinal cortex (MEC) is home to several functional classes of neurons that encode information relevant for spatial navigation. For instance, border cells [1] are neurons that are active whenever an animal is near a boundary, suggesting they represent “affordances” (i.e., features of an environment that shape how navigation is constrained). Similarly, grid cells [2, 3] are neurons that are active along vertices of a triangular lattice in space, enabling them to represent a coordinate system, which could be useful for encoding trajectories through space. This perspective has led to hypotheses that grid cells play an integral role in path-integration [4], a cognitive process that involves updating an estimate of where in space an organism is from a sequence of movement directions and speeds.

Experimental evidence for the brain’s use of grid cells in path-integration has been challenging to obtain, as it requires both quantitatively measuring path-integration in animals and inactivating/manipulating grid cells. However, there exist some results supporting this hypothesized role [5]. Additional evidence that grid cells are involved in the neural computations underlying path-integration comes from work that trained recurrent neural networks (RNNs) to perform path-integration and found grid-like representations emerging in the RNN’s recurrent layer [6–8]. This has provided a “normative” explanation for grid cells. Namely, that they are general solutions to the problem of path-integration. While the biological plausibility of these RNN models remains debated [9–11], comparison of trained model activations and real neural data has shown that these RNNs can capture non-trivial properties of grid cell and MEC activity [12, 13].

The putative role of MEC, and grid cells specifically, in path-integration has suggested that they may additionally be shaped by predictive information [14, 15]. Such prospective coding could be leveraged for navigational planning [16–18] and composition [19]. In-line with this hypothesis, neurophysiological recordings have found grid cells have an optimal shift forwards in time, with grid score – which measures the extent of the hexagonal grid-like structure – increasing when considering not the animal’s current position, but its future position [20, 21]. Furthermore, stationary rodents exhibit “theta-sweeps” in MEC and hippocampus, which includes the activity of grid cells at future locations in space [22–25]. Recently, a new functional class in MEC was discovered that has grid structure only when neural activity is projected onto future spatial locations, making them a distinct subpopulation from standard grid cells [26]. While on the face of it these “predictive grid cells” appear consistent with previous work demonstrating prospective coding in MEC, several questions remain. In particular, if existing theories already attribute computational roles in predictive coding to standard grid cells [14–19], how should predictive grid cells be understood?

Here, we aim to probe features of predictive grid cells that may provide insight into their computational role. First, we re-analyze state-of-the-art MEC recordings [27] that were done in a facility and with an experimental environment that are distinct from those used in the original discovery of predictive grid cells [26]. If predictive grid cells really are a fundamental component of the MEC’s involvement in cognitive processes, then we expect signatures of predictive grid coding to also be present in this separate dataset. In-line with this, we find MEC cells that are not standard grid cells, but have strong grid scores when considering the future spatial location of the rodent. We then examine whether predictive grid coding emerges in RNNs trained to perform path-integration [6–8]. If predictive grid cells are used for goal-directed route planning [26], then the RNNs should not develop such a representation, as the trajectories the RNNs are trained to path-integrate are short and pseudo-random. In contrast, if predictive grid cells are involved in maintaining and updating the grid cell network (e.g., through the continuous attractor network (CAN) that is believed to underlie standard grid cells [28]), then path-integrating RNNs should develop predictive grid coding. Consistent with this second hypothesis, we find recurrent units in the RNNs that exhibit predictive grid coding. We show that this is a robust feature, with similar percentages of predictive grid units present in 10 independently trained path-integrating RNNs. Furthermore, we find that the recurrent units classified as predictive grid units are significantly more sensitive to the movement direction than standard grid units, providing strong evidence that the identified units are truly encoding prospective information. Recurrent units that exhibit band-like tuning, analogous to band cells in MEC [29], have been argued to play an important role in path-integrating RNNs [7, 30–33]. We find that these band units non-trivially overlap with predictive grid units, illustrating that predictive grid coding may be broadly present in the RNNs, especially in units involved in updating grid unit activity. Finally, we return to the MEC electrophysiology recordings and look for a signature of predictive grid cell coding during sleep. While limited, we find one recording where predictive grid cells – identified during open field exploration – exhibit a bias towards encoding information about the toroidal manifold the population activity lies along on [27] with prospective, but not retrospective shifts.

Our work demonstrates that predictive grid coding is a general feature of rodent MEC and that explicit planning is not needed for such representations to emerge in path-integrating RNNs. Instead, we hypothesize that predictive grid cells play a role in maintaining and updating the grid cell population dynamics, a prediction that can guide future experimental and theoretical analysis.

## Results

### Predictive grid cells are a general feature of rodent MEC

We begin by re-analyzing previously recorded and published large-scale electrophysiology recordings of rat MEC [27]. While this dataset contains a large number of grid cells, which preclude them from being classified as predictive grid cells [26], these recordings represent the state-of-the-art and achieved simultaneous recording of multiple grid modules. Therefore, we reasoned that it would be a good dataset to search for predictive grid cells in.

For each recorded cell (*N* = 1330), we compute its grid score. We classify those with grid score above a threshold value of 0.5 as standard grid cells (766 of the 1330 cells passed this criteria). For all the remaining cells (564 cells), we shift the recorded spike times prospectively (Δ *>* 0 ms) and retrospectively (Δ *<* 0 ms), with respect to the position of the rats, finding the shift that achieves the maximal grid score, Δ_max_. We then define predictive grid cells as those cells whose Δ_max_ *>* 0 ms and whose grid score at Δ_max_ is above a threshold of 0.3. To ensure that this threshold is sufficiently strong, we perform a shuffle analysis for each cell, generating a distribution of grid scores that would be expected by chance. We further require that the grid score at Δ_max_ for all putative predictive grid cells is above the 95^th^ percentile (see Methods, for more details).

Across all 9 recordings (*n* = 3 rats), we find 117 cells that pass the criteria for being classified as predictive grid cells (median % of non-grid cells classified as predictive grid cells, per recording: 22.4%; range: 4% *−* 52%). We present the results from a single module and rat (recording ID R21) in Fig. 1. Additional results from other rats and modules can be found in Fig. S1. Example cells that are identified as predictive grid cells are shown in Fig. 1A. In each case, despite having a low grid score at Δ = 0 ms, prospective shifts in time lead the associated ratemaps and spatial autocorrelograms (SACs) to become increasingly grid-like (Fig. 1A, B). This increase is selective to shifts around 100 *−* 200 ms (Fig. 1C), as larger shifts lead to a decrease in grid score (Fig. 1B). These results are largely consistent with what the original work that discovered predictive grid cells found when shifting in time [26]. In addition, when we perform shifts in space by projecting along the estimated movement direction of the rats, we again find cells that exhibit increased grid scores with prospective shift (Fig. S2). Maximum grid score and percent of identified predictive grid cells were comparable between temporal and spatial shifts (Fig. S3).

**Figure 1:**
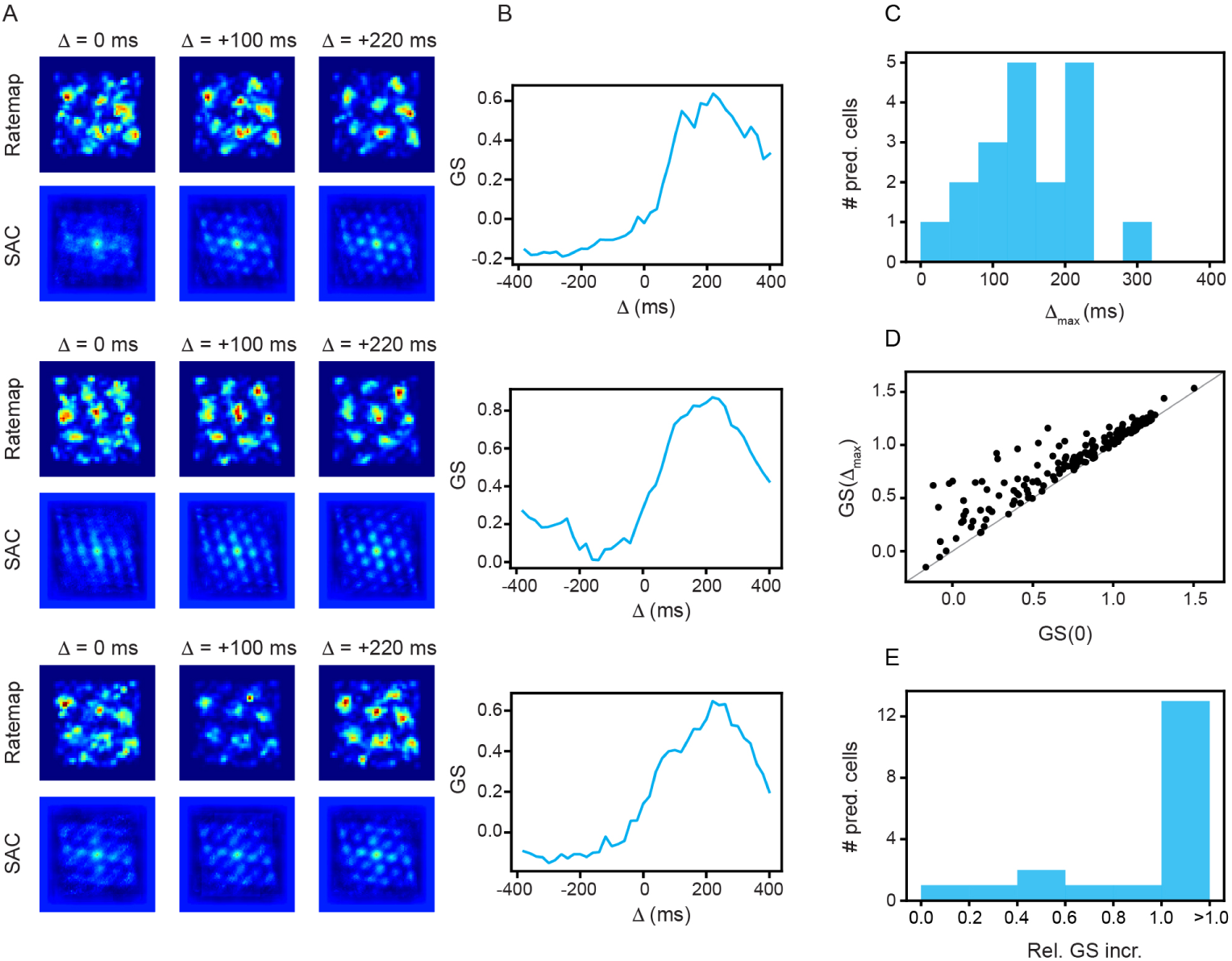
Predictive grid cells are present in Gardner et al. (2022) MEC recordings (recording ID R21). (A) Example ratemaps and spatial autocorrelograms (SACs), at different temporal shifts, for cells classified as predictive grid cells. (B) Grid score, as a function of temporal shift, tuning curves for the three predictive grid cells in (A). (C) Distribution of temporal shifts with maximal grid score (Δ_max_) for all cells classified as predictive grid cells. (D) Grid score at Δ = 0 ms, compared to grid score at Δ = Δ_max_, for all recorded cells. (E) Distribution of relative grid score increase for all cells classified as predictive grid cells.

The increase in grid score with prospective temporal shifts is largely confined to cells with small grid score when no shift (Δ = 0 ms) is applied (Fig. 1D). For those cells with a high grid score at Δ = 0 ms, little benefit is observed from shifting in time, indicating that this increase occurs in a specific subset of cells. To quantify the improvement in grid score we observe when shifting prospectively in time, we compute the relative grid score increase, which is given by <u>^GS(Δmax)^</u>*<u>^−^</u>*<u>^GS(0)^</u> , where GS(Δ) denotes the grid score at a Δ ms shift. If GS(Δ_max_) *>>* GS(0), the relative grid score will be *>* 1. We find that the distribution of relative grid scores for all cells classified as predictive grid cells is skewed towards values *>* 1 (Fig. 1E), demonstrating that the majority of the cells we classify as predictive grid cells do not just meet the criteria, but see a large increase in grid score when shifted prospectively, providing further support of their predictive grid tuning.

We also identify some recorded cells as having their Δ_max_ *<* 0, making them tuned towards retrospective shifts (Fig. S4A). However, examining their grid score as a function of Δ curves, we find that the tuning is noisy and weak (Fig. S4B), which is further supported by the distribution of relative grid score increases being primarily restricted to *<* 0.5 (Fig. S4D).

Taken together, our results (done on a separate dataset from the original work of Ouchi and Fujisawa (2024) [26], using separate analysis code) provide evidence that predictive grid cells are a general feature of rodent MEC.

### Predictive and retrospective grid units emerge in path-integrating RNNs

Having established that predictive grid cells are present in an independent set of MEC recordings, we next ask whether a comparable representation emerges from the computations required for path-integration. We trained RNNs using same the architecture and hyperparameters of previous path-integrating RNN models [6, 8]. The networks navigated a 2.2 *×* 2.2 m^2^ environment along pseudo-random trajectories, each lasting 800 ms. At the beginning of each trajectory, the network received a learned encoding of its initial allocentric position, and then updated its position estimate using the agent’s velocity at each time step (see Methods, for more details). These networks provide a useful setting in which to examine the origin of predictive grid coding. Their objective requires continuous position updating, but the actual training task has no goal selection or route planning component. Therefore, predictive grid cell representations that emerge under these conditions can be attributed to the recurrent solution that is learned for path-integration.

We analyze the spatial tuning of the recurrent units using the same temporal shifting approach that was applied to the MEC recordings (see Methods). We find individual units exhibit a diversity in how temporal shifting alters the spatial structure of their activity (Fig. 2A). For instance, a unit that was classified as a standard grid unit, with a clear hexagonal pattern in its SAC when its activity was aligned with the agent’s current position (Δ = 0 ms) (Fig. 2A, top), shows an approximately symmetric decrease in grid score with increasing retrospective and prospective shifts (Fig. 2B, top). In contrast, a unit that was classified as a predictive grid unit had weak grid structure at Δ = 0 ms, but exhibited a monotonic increase in grid score until Δ = 240 ms (Fig. 2A, B, middle). Similarly, a unit that is classified as retrospective grid unit demonstrates a large increase in grid score for retrospective shifts (Δ *<* 0 ms; Fig. 2A, B, bottom).

**Figure 2:**
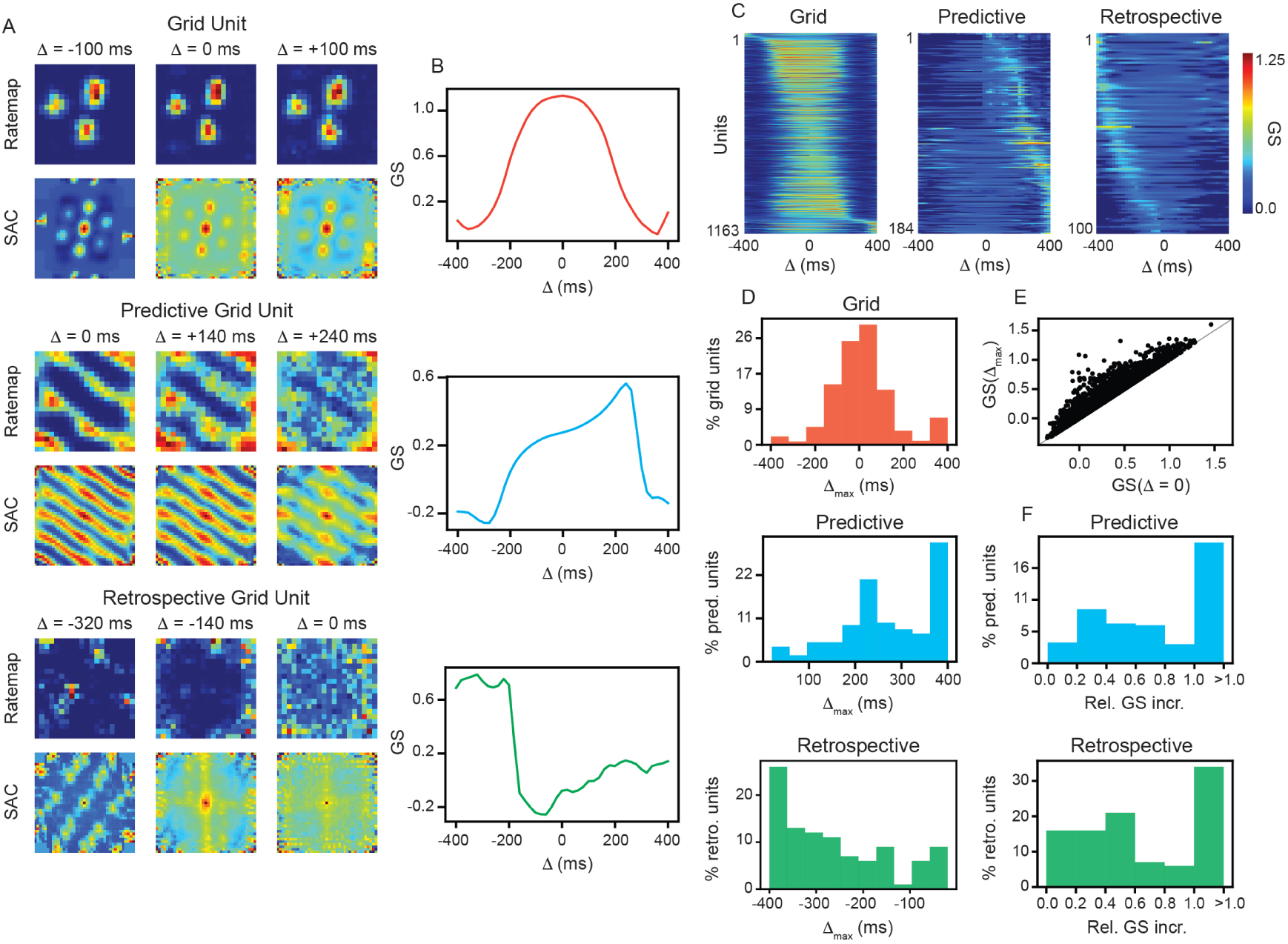
Predictive and retrospective grid units are present in RNNs trained to perform path-integration. (A) Example ratemaps and SACs, at different temporal shifts, for units classified as grid, predictive grid, and retrospective grid units, respectively. (B) Grid score, as a function of temporal shift, tuning curves for the three predictive grid cells in (A). (C) Grid score, as a function of shift, for all classified, grid, predictive grid, and retrospective grid units, respectively. (D) Distribution of Δ_max_ for all classified, grid, predictive grid, and retrospective grid units, respectively. (E) Grid score at Δ_max_ compared to grid score at Δ = 0 ms for all non-dead RNN units. *x* = *y* line denotes units with Δ_max_ = 0 ms. (F) Distribution of relative grid score increase for classified predictive grid and retrospective grid units, respectively.

The distinction between these example units is apparent across the entire recurrent population (Fig. 2C), with units classified as standard grid units generally maintaining the highest grid scores near Δ = 0 ms and predictive (retrospective) grid units having their highest grid scores after their activity was shifted towards future (past) positions. In the RNN analyzed in Fig. 2, out of 3080 units that exhibited sufficient activity (“non-dead units”), we find 1163 standard grid units (37.8%), 184 predictive grid units (6.0%), and 100 retrospective units (3.3%). While the population of standard grid units have their Δ_max_ clustered around 0 ms, predictive and retrospective grid units exhibit a broad distribution of Δ_max_ (Fig. 2D). As with the MEC data (Fig. 1D, E), we find that many recurrent units exhibit a large increase in grid score with temporal shift (Fig. 2E) and that the relative grid score increase of both predictive and retrospective units is strongly skewed towards value *>* 1.0 (Fig. 2F). We also find predictive and retrospective grid units when shifting in space, instead of in time (Fig. S5), with similar relative grid score increases and percent units identified as predictive grid units as was found with temporal shifting (Fig. S6).

Having identified predictive and retrospective grid units in an individual path-integrating RNN, we then examine whether they would reliably emerge across multiple training runs. We therefore analyze 10 independently initialized RNNs that were trained on path-integration. Across all networks, we find predictive and retrospective grid units populations (Fig. 3A). On average, predictive units represented 4.7% (range: 3.4% *−* 6.0%) of non-dead units, while retrospective units represented 3.4% (range: 2.2% *−* 4.8%). We note that, while this percent of classified predictive and retrospective grid units is considerably lower than the percent of standard grid units (median: 38.2%; range: 32.4% *−* 42.4%), we use particularly conservative thresholds for what is classified as a predictive and retrospective grid unit, and particularly liberal thresholds for what is classified as a standard grid unit (see Methods)^1^. Consistent with what we found when looking at a single RNN, we find that the predictive and retrospective grid units exhibit a large increase in relative grid score (Fig. 3B) and, unlike standard grid units, have a broad distribution of Δ_max_ (Fig. 3C).

**Figure 3:**
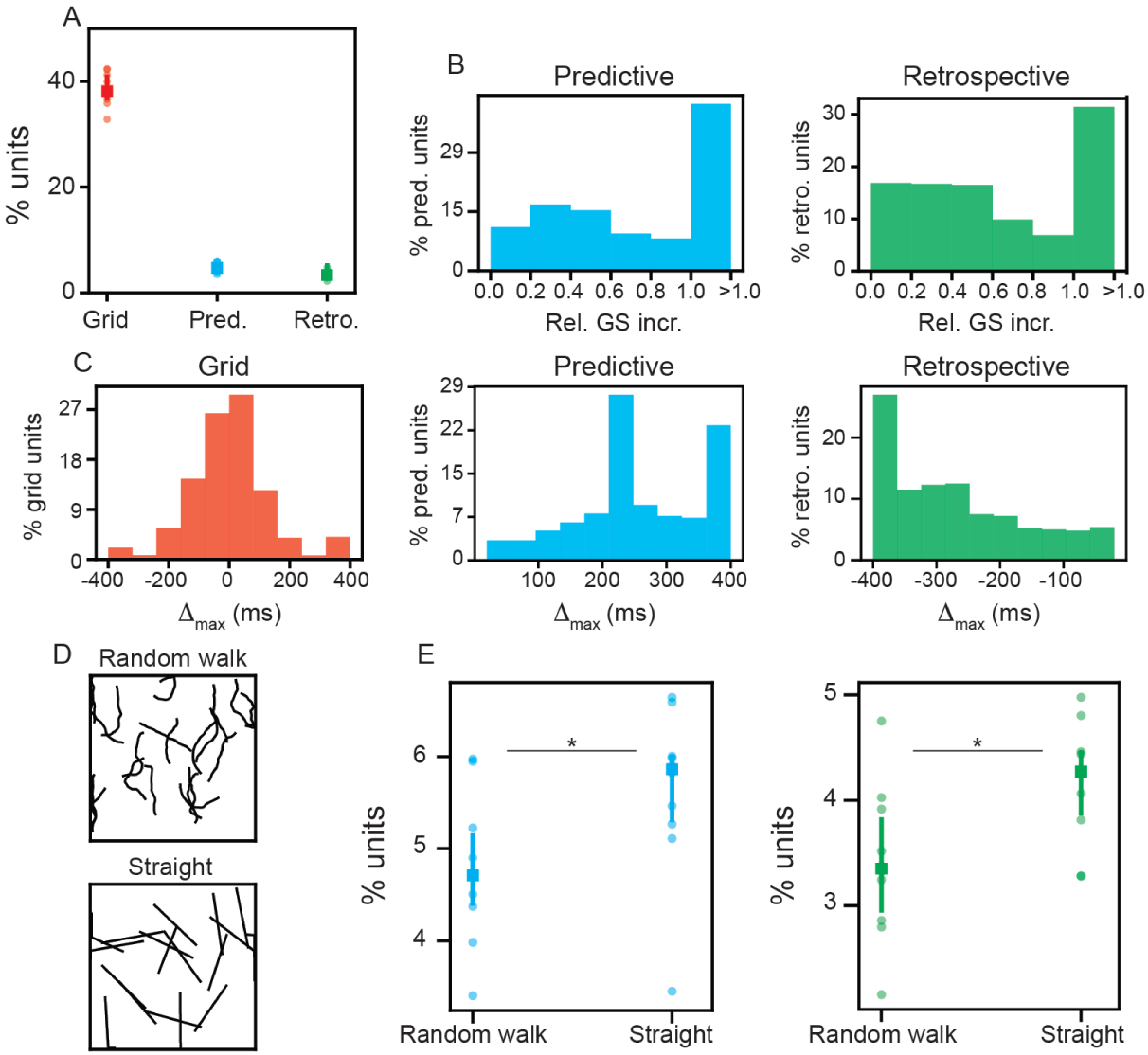
Predictive and retrospective grid coding robustly emerges across independently trained path-integrating RNNs. (A) Percent of all non-dead RNN units classified as grid, predictive grid, and retrospective grid units, across 10 independently trained networks. (B) Distribution of relative grid score increase for classified predictive grid and retrospective grid units, across all 10 independently trained networks. (C) Distribution of Δ_max_ for all classified, grid, predictive grid, and retrospective grid units across all 10 independently trained networks. (D) Illustration of example random walk and straight path trajectories sampled to generate ratemaps. (E) Percent of classified predictive and retrospective grid units using random walk and straight path trajectories. Each dot represents one network, square denotes median, and error bars denote 25^th^ and 75^th^ percentiles across all 10 independently trained networks. Star denotes *p*-value *<* 0.05 using one-side Kolmogorov-Smirnov (KS) test.

A major finding by Ouchi and Fujisawa (2024) [26] was that predictive grid coding was increased when rodents ran along paths with stereotyped structure (e.g., straight paths) than when rodents ran along less constrained routes (e.g., pseudo-random trajectories). Therefore, we examined whether the RNNs, trained to path-integrate pseudo-random trajectories, exhibited more predictive grid coding when then evaluated on straight paths (Fig. 3D). We find a significant increase in units classified as predictive and retrospective grid units on these straight path trajectories (Fig. 3E; median % classified as predictive: 5.9%; median % classified as retrospective: 4.3%; *p−*value of one-sided Kolmogorov-Smirnov (KS) test predictive: 0.026; *p−*value of one-sided KS test retrospective: 0.026). We do not find a significant increase in the percent of units classified as standard grid units (median: 39.5%; *p−*value of one-sided KS test: 0.21).

Another important feature of predictive grid cells is that they are future path-dependent [26]. That is, the activity of a predictive grid cell is dependent not only on the allocentric location, but also on the movement direction of the rodent. Thus, if the recurrent units classified as predictive grid units in the path-integrating RNNs are truly encoding prospective information, we expect that their activity will similarly be future path-dependent. To test this, we bin the environment into 0.05 *×* 0.05 m^2^ bins and compute the population vector correlation of predictive grid units for pairs of trajectories that intersect the same spatial bin at the same time-step along the trajectory (Fig. 4A; see Methods, for more details). We also compute the population vector correlation of grid units, to enable a comparison. We find that predictive grid units exhibit a significantly broader distribution of population vector correlations than grid units (Fig. 4B). In addition, we find that the population vector correlation of predictive grid units is significantly more dependent on the

**Figure 4:**
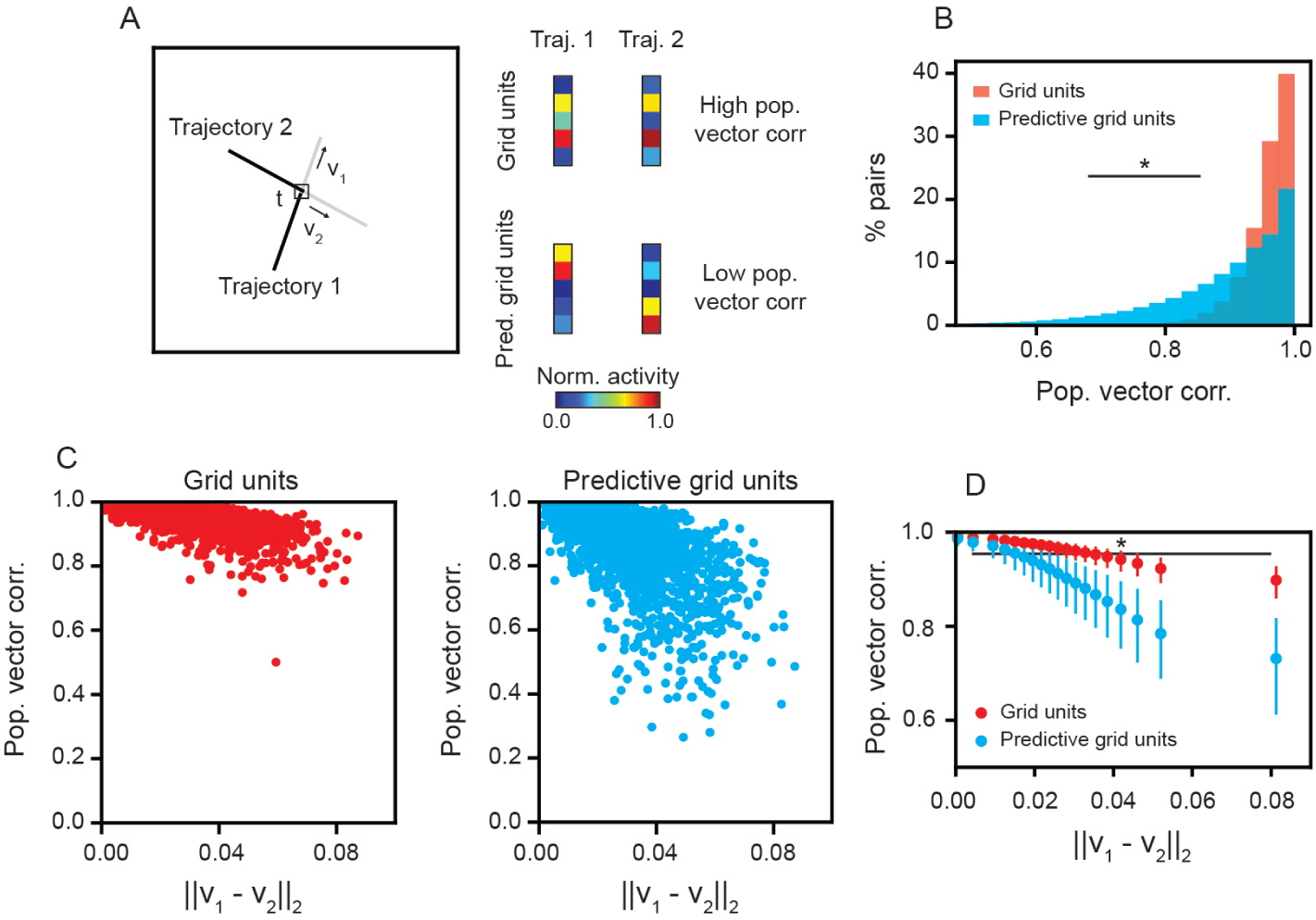
Population vector correlation is more future path-dependent for predictive grid units than grid units. (A) Left – Schematic illustration of two trajectories that go through the same spatial bin of the environment at time-step *t*. Black line denotes the trajectories up to time-step *t* and gray line denotes the future paths after time-step *t*. The movement vectors of the two trajectories are denoted by *v*_1_ and *v*_2_. Right – Schematic illustration of possible values of the grid and predictive grid population vectors at time-step *t*, for each trajectory. Grid units are hypothesized to have a higher, less future path-dependent, population vector correlation than the predictive grid units. (B) Distribution of population vector correlations at pairs of trajectories that intersect at time-step *t* = 20, for grid and predictive grid units. (C) Population vector correlations, for grid and predictive grid units, as a function of Euclidean distance between the movement vectors of the two trajectories, *||v*_1_ *− v*_2_*||*_2_. For visualization purposes, every 100^th^ pair of trajectories is plotted. (D) Average population vector correlation for binned Euclidean distance between the movement vectors of the two trajectories. Bins were chosen as the 5^th^, 10^th^*, …,* 95^th^ percentiles. Dots denote median value and lines denote 25^th^ and 75^th^ percentiles computed across all pairs of trajectories. In (B) and (D), star denotes *p*-value *<* 0.05 using one-sided KS test.

Euclidean distance between the movement directions of the two trajectories than grid units (Fig. 4C, D). That is, when the two trajectories are aligned and moving in a similar direction (*||v*_1_ *− v*_2_*||*_2_ *≈* 0), then the predictive grid unit population vectors are highly correlated. However, when the two trajectories are not aligned (*||v*_1_ *− v*_2_*||*_2_ *>* 0), despite having a similar allocentric location, the predictive grid unit population vectors are weakly correlated. Similar results are found when choosing different values of time in which the trajectories intersect (Fig. S7). Thus, while the recurrent units classified as standard grid units are largely independent of the future path, and therefore, show strong allocentric coding, those classified as predictive grid units are significantly more future path-dependent. This provides further evidence that path-integrating RNNs develop units with predictive grid coding.

These results demonstrate that predictive and retrospective grid coding can emerge in recurrent networks trained solely to update position from self-motion, even when the training objective contains no explicit demand for planning.

### Band units exhibit predictive grid coding

Path-integrating RNNs are known to develop units with spatial tuning beyond grid-like patterns. In particular, band-like representations [7] and border-like representations [6, 7] have been found to emerge. This tuning is analogous to band cells [29] and border cells [1], functional classes of neurons found in MEC. Detailed examination has shown that band and border units in RNNs play integral roles in path-integration [7, 30–33], demonstrating their importance in the computations learned by the RNN.

Given that examples of predictive and retrospective grid units appear to have band-like and border-like responses at Δ = 0 ms (Fig. 2A), we examine whether border and/or band units exhibit stronger predictive or retrospective grid coding. As in previous analysis, we find units in the RNN with high band and border scores (Fig. 5A, B; see Methods, for more details). We classify units as band or border units by taking the 95^th^ percentile of the band and border score distributions (for each independently trained network), respectively. We then compute the percent of classified border and band units that are also classified as grid, predictive grid, or retrospective grid units. We do not see significantly greater overlap with grid units than expected by chance (Fig. 5C, left). This is due to the fact that, in order for a unit to be classified as a grid cell, it must have separable peaks in its SAC (Fig. 2A), which is less likely to happen when the underlying ratemap has strong band-like or border-like responses. In contrast, for both predictive and retrospective grid units, there is a significantly greater overlap with classified band units than chance (Fig. 5C, middle and right). This is unique to band units, as neither predictive nor retrospective grid units overlap significantly with border units.

**Figure 5:**
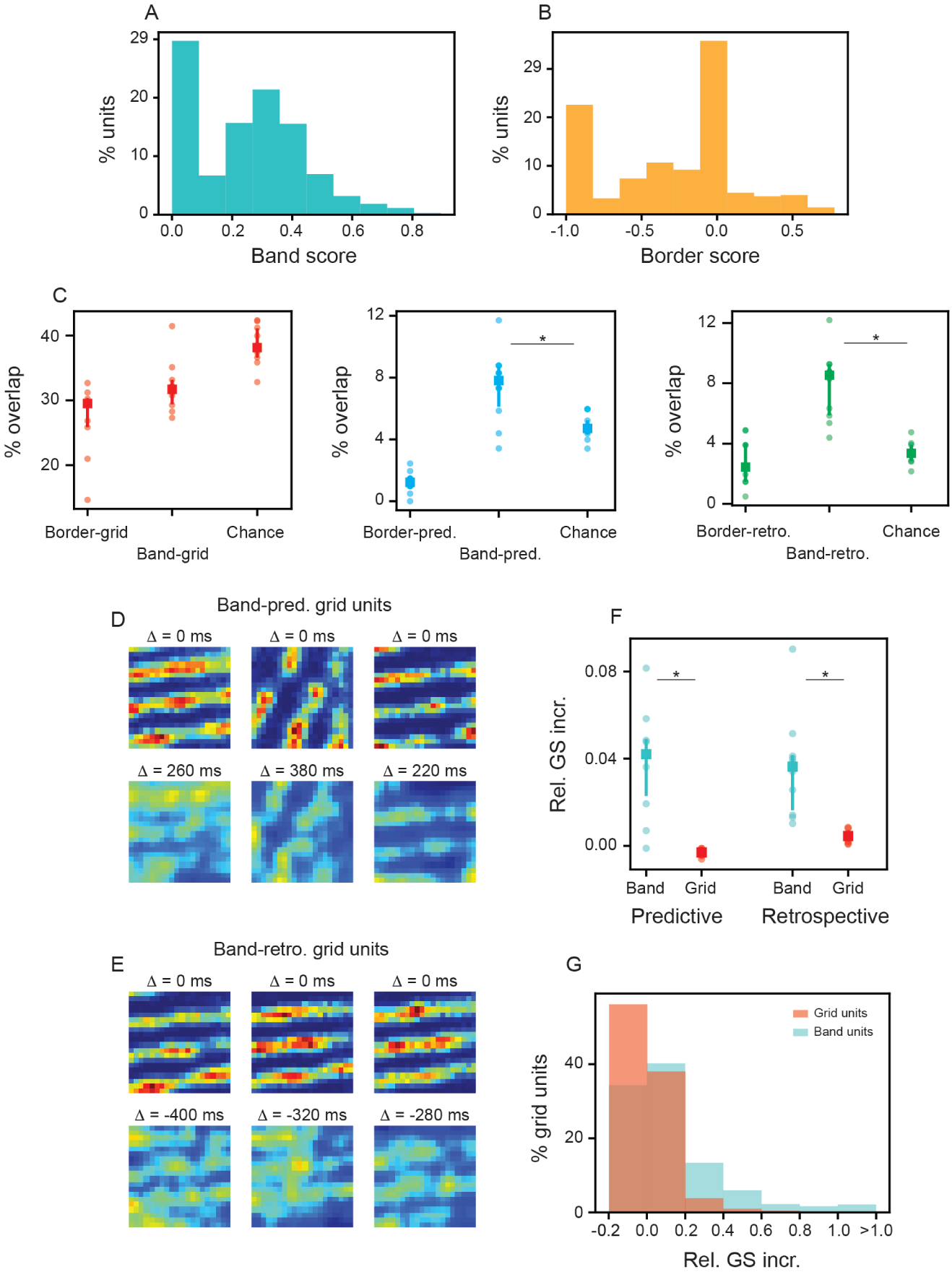
Band units exhibit predictive and retrospective grid coding. (A) Distribution of band scores, across all units in 10 independently trained networks. (B) Distribution of border scores, across all units in 10 independently trained networks. (C) Percent of units classified as border or band units and grid, predictive grid, or retrospective grid units, respectively. Each dot corresponds to the value of an individual network. Squares denote median value and bars denote 25^th^ and 75^th^ values across all 10 independently trained networks. (D)–(E) Three example ratemaps at Δ = 0 ms and Δ_max_ for units that are classified as both band and predictive grid or band and retrospective units, respectively. (F) Relative grid score increase for band and standard grid units when shifting prospectively or retrospectively. Each dot corresponds to the value of an individual network. Squares denote median value and bars denote 25^th^ and 75^th^ values across all 10 independently trained networks. (G) Distribution of relative grid score increase for all band and standard grid units. In (C) and (F), Star denotes *p*-value *<* 0.05 using one-sided KS test.

We visualize example units that are classified as both band and predictive grid units (Fig. 5D), and units that are classified as both band and retrospective grid units (Fig. 5E). While the activity is noisy at Δ_max_, grid structure – including structure that is distinct from that present at Δ = 0 ms – can be seen. This demonstrates that the units in the overlap contain reasonable encoding of both band and predictive or retrospective grid tuning. If band units broadly exhibit predictive and retrospective grid coding, then we expect large relative increases in grid score when shifting either prospectively or retrospectively, even for units that are not classified as predictive or retrospective grid units. In-line with this hypothesis, we find an average relative increase (for each individually trained RNN) in grid score for band units that is significantly greater than that found for grid units, when shifting either prospectively or retrospectively (Fig. 5F). This difference also holds at the individual unit level (Fig. 5G), where 37.5% of band units exhibit a relative grid score increase of *>* 10% when shifting prospectively, compared to only 11.9% of grid units. We also find that band units have a significantly greater relative increase of grid score than border units (Fig. S8). These results provide evidence that band units, as a function class generally, contain predictive and retrospective grid coding, suggesting that such coding may be linked to their role in updating spatial representations during path-integration.

### Predictive grid cell signature is present during sleep

Given our finding that predictive grid units emerge in path-integrating RNNs, even in the absence of a predictive objective, and that predictive grid coding is present in band-units, which have been found to play a major role in path-integration in RNNs [30, 33], we hypothesize that predictive grid cells may be a property of the grid cell circuit. Previous work has shown that grid cells maintain their between-cell relationships during sleep [34, 35] and that left-right-alternating theta-sweeps are present in MEC during sleep [24]. Guided by our hypothesis, we test whether a signature of predictive grid coding is present during sleep.

Inspired by recent work showing that grid cells exhibit tuning to specific locations on a two-dimensional torus that can be extracted from grid cell population activity [27] (“toroidal ratemaps”, Fig. 6A; see Methods, for more details), we characterize how that tuning changes with prospective and retrospective temporal shifts. During exploration of an open field (OF) (the same recording as was analyzed in Fig. 1), we find that predictive grid cells appear to maintain their tuning to toroidal coordinates better when shifting prospectively than retrospectively (Fig. 6B, left). In contrast, for cells that were not classified as predictive grid cells (“non-predictive grid cells”), we observe little difference when shifting prospectively or retrospectively (Fig. 6B, right). We quantify this qualitative finding by computing the spatial information [36] present in the toroidal ratemaps (see Methods). We find that for predictive grid cells, but not for non-predictive grid cells, spatial information stays significantly higher when shifting prospectively, as compared to retrospectively (Fig. 6C, D). This same trend is found in four of the eight recordings we analyze^2^ (Fig. S10: R11, R12, Q1) and in another recording it trends towards significance (Fig. S10: R22).

**Figure 6:**
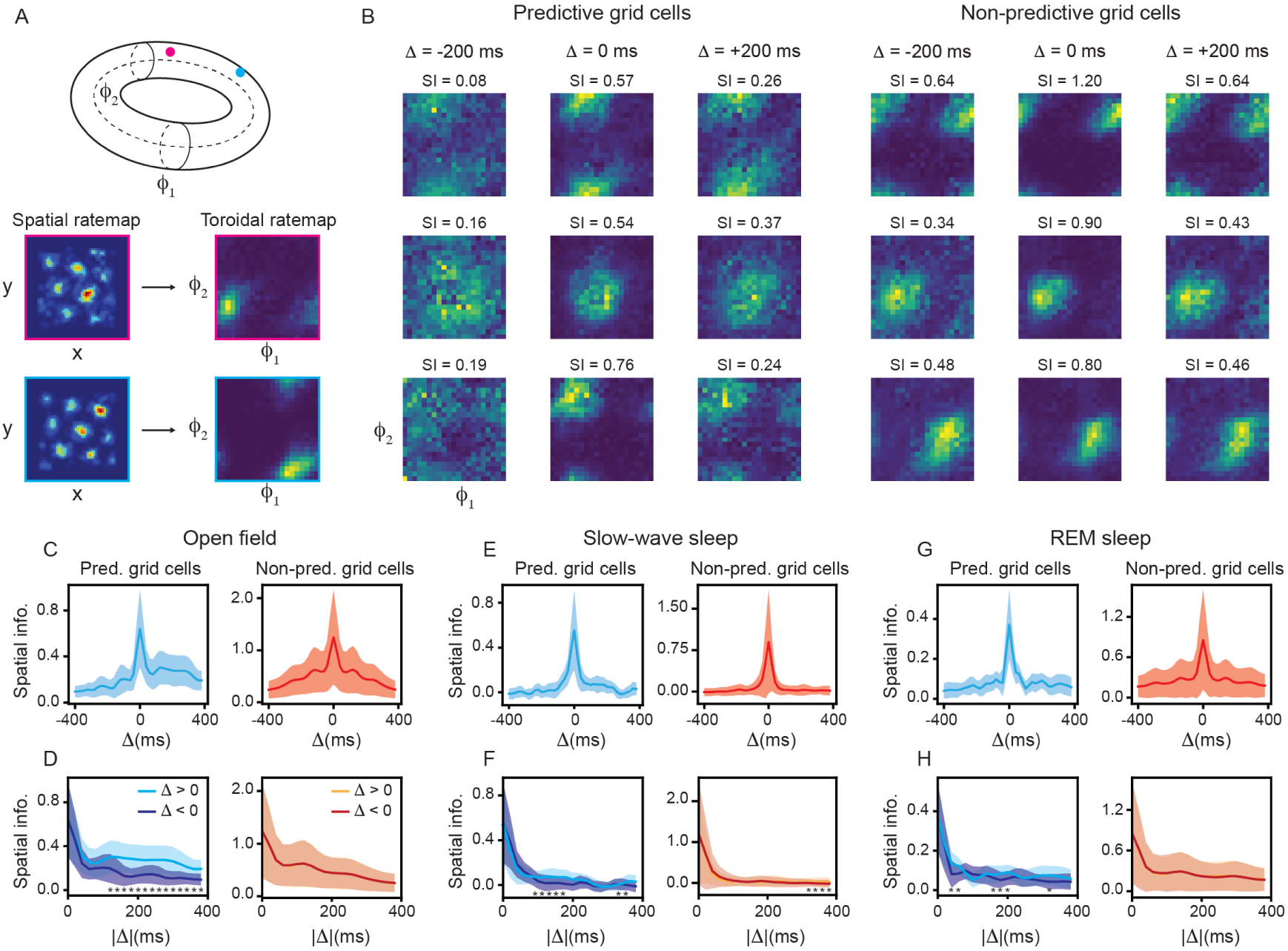
Predictive grid cells exhibit greater selectivity in their toroidal ratemaps when shifting prospectively, including during sleep (recording ID R21). (A) Schematic illustration of toroidal ratemaps. (B) Example toroidal ratemaps, constructed using three different temporal shifts, from three cells identified (left) and not identified (right) as predictive grid cells. (C) Spatial information of toroidal ratemaps, as a function of temporal shift, for predictive and non-predictive grid cells, recorded in the open field. (D) Same as (C), but with retrospective and prospective shifts plotted on the same axis. (E)–(F), (G)–(H) Same as (C)–(D), but for recordings during slow-wave and REM sleep, respectively.

We then perform the same analysis on data collected while the same rats were asleep. Because these recordings were done all during the same experimental sessions, we could be reasonably confident that extracted neuron labels, through spike-sorting [37], were consistent across sleep and waking. This means that the predictive grid cells we identify in the OF condition can be used to study how predictive grid cells behave during sleep. Comparing average spiking rates of predictive and non-predictive grid cells, we find no statistically significant difference when considering OF exploration, slow-wave sleep, or REM sleep (Fig. S9), suggesting that predictive grid cells are just as active as standard grid cells during sleep. Of the four recordings with predictive grid cells that exhibited toroidal ratemaps with significantly greater spatial information when shifting prospectively than retrospectively, only two had associated sleep data^3^. In one of these two recordings, we find that predictive grid cells again exhibited significantly greater spatial information for some prospective shifts (Fig. 6E–H; see Fig. 6 for other recording), both when considering slow-wave and REM sleep. We discuss possible explanations for this limited observation in the Discussion. Our results demonstrate that a non-trivial prediction of our path-integrating RNN analysis, namely that predictive grid cells should be present even in the absence of explicit planning, is supported by one experimental recording of predictive grid cell activity during sleep.

## Discussion

Grid cells have historically been believed to support path-integration by maintaining a spatial representation that is continuously updated from self-motion [4]. As a growing number of experiments have shown that medial entorhinal cortex exhibits similar representations in non-spatial (“abstract”) spaces [38–40] and that rewards can affect these representations [41–43], there has become a growing appreciation that grid cells are also involved in general goal-directed navigation. Because a key aspect of this cognitive process involves prediction (e.g., predicting when a reward is coming), predictive coding has become another lens with which to understand grid cells [14–19]. The discovery that MEC also contains cells that encode the animal’s prospective position, in the form of a “predictive grid code” [26], has been viewed as further supporting the predictive coding perspective of grid cells. However, given that existing predictive coding theories have explained aspects of grid cells (and not required any prospective shifting), it has remained unclear what functional role predictive grid cells play. Previous computational attempts to study predictive grid cells have largely left this question open, studying instead the biophysical mechanism that could enable predictive grid tuning [44] or imposing a predictive objective on the model [45]. In addition, because predictive grid cells have been found in only a single dataset, it remains unclear how general a feature such coding even is in MEC.

To address this, we performed analysis of both MEC electrophysiology recordings and path-integrating recurrent neural networks. We found clear signatures of predictive grid coding in a state-of-the-art MEC dataset (Fig. 1), which had been collected in a different facility, under different experimental conditions, using different analysis code as the original work describing predictive grid cells. We found predictive grid cells when projecting prospectively from the animal’s current position using either space or time (Figs. 1, S1 – S3). In addition, consistent with Ouchi and Fujisawa (2024) [26], we found that predictive grid cells exhibit a range of prospective shifts over which their maximal grid score is achieved, *≈* 300 ms for temporal shifts and *≈* 0.15 m for spatial shifts. We also find predictive grid cells demonstrate a bias towards prospective shifts when considering how their activity encodes the toroidal coordinates of the underlying population manifold (Fig. 6B–D), a new analysis that was not previously performed. These results provide evidence that predictive grid coding is a general feature of rodent MEC.

We found similar evidence of predictive grid coding in path-integrating RNNs (Fig. 2). While the biological plausibility of these networks remains debated [9–11], the emergence of predictive grid coding in the RNNs provides an unbiased test of alignment with MEC coding, further supporting previous claims of consistency between the networks and neural data [12, 13]. The predictive grid units we identified exhibited large increases in grid score at their maximal temporal shift (Fig. 3B) and had significantly greater path-dependence than standard grid units (Fig. 4). In addition, we found a significant increase in the number of units classified as predictive grid units when evaluating the networks on straight paths, as compared to pseudo-random walks (Fig. 3E), consistent with the observation that predictive grid coding was more prevalent when rodents ran along stereotyped trajectories, as opposed to freely explored [26]. Collectively, these results demonstrate that the predictive grid coding we found in the path-integrating RNNs is not merely an artifact of our selection criteria or driven by noisy responses [46]. Therefore, we posit that predictive grid coding does not require explicit planning to emerge.

We examined whether border or band units – which have previously been shown to play an important role in the networks [7, 30, 32, 33] – showed increased grid scores when shifting their prospectively in time. We found band units, but not border units, exhibited more overlap with predictive grid units than expected by chance and experienced a larger relative increase in grid score when shifting prospectively (Fig. 5). In addition to having implications for our understanding of the role predictive grid coding plays in path-integrating RNNs, we hypothesize that this may also resolve a discrepancy between such networks and MEC recordings. Namely, if band units in the RNNs are functionally playing a similar role as predictive grid cells in MEC, then this could explain why band units have been consistently reported in path-integrating RNNs [6, 7, 30, 32, 33], while they have been largely absent from neurophysiological data (with the exception of the original discovery of band cells [29]).

Previous work has shown that left-right-alternating theta-sweeps, which have been viewed as evidence of predictive planning by the hippocampus [22, 23], has recently been shown to be driven by left-right-alternating theta-sweeps in MEC grid cells [24, 47], which are present during sleep [24, 25] and in parts of arenas that do not allow for any choices in movement [47] (i.e., have no planning component). This suggests that this “predictive” signal may be driven by local circuits in MEC [24], such as those involved in maintaining continuous attractor dynamics [28]. Because we found predictive grid units in path-integrating RNNs, we hypothesized that predictive grid cells might similarly be a features of the underlying circuit, as opposed to a direct consequence of planning. To test this, we looked for a signature of predictive grid coding during sleep. Namely, a bias in coding toroidal coordinates when shifting prospectively, as compared to when shifting retrospectively. We found this signature in one of the two recordings that had MEC activity recorded during sleep and exhibited the prospective bias during open field exploration. This limited observation should be interpreted cautiously, and may be due to that specific recording having the longest amount of continuous sleep data and/or the topological data analysis (TDA) methods better capturing the underlying toroidal structure. Indeed, the TDA analysis pipeline has several different hyperparameters which can lead to sharper resolution of the torus, but we left these values as they were to be consistent with the previously reported results [27]. It may be that better choices could lead to greater identification of predictive grid cell coding during sleep. Furthermore, the toroidal coordinates are estimated using both predictive and non-predictive grid cell activity. An important determinant may therefore be the ratio of predictive to non-predictive grid cells, as the more evenly matched the two are the more predictive grid cells influence the estimated torus. Given the challenges related to numerically approximating the underlying toroidal manifold, we view the fact that we found any recording where predictive grid cell coding is present during sleep as a non-trivial result, particularly as this is not an obvious prediction from the existing explanation of predictive grid cells [26, 45].

### Limitations

Because a cell must exhibit a low grid score when its activity is aligned with the animal’s current position in order to be classified as a predictive grid cell [26], standard grid cells and predictive grid cells must necessarily form distinct subpopulations of MEC. However, the neurophysiological dataset we analyzed was post-processed before being made publicly available to only include putative grid cells [27] (see Methods). This means that the data was strongly biased, with over half of the cells (58%) being too grid-like to be considered as candidates for predictive grid cells. Therefore, we view our analysis as not necessarily reflecting the true distribution and prevalence of predictive grid coding in MEC. Despite this, we found evidence of predictive grid cells, and the measured preference for temporal and spatial shifts were mostly consistent with what was reported by Ouchi and Fujisawa (2024) [26].

While we find predictive grid units in all 10 independently trained RNNs (Fig. 3), there is also variability in the extent to which they overlap with band and border units (Fig. 5). This may be due to “solution degeneracy” in RNNs [48]. Increasing the number of independently trained RNNs may enable better assessment of statistical significance. In addition, restricting ourselves to analyzing specifically those networks that are most dependent on predictive grid cells may lead to a better understanding of what role predictive grid cells can play.

More generally, our analysis focused on binary classification, with MEC cells or RNN units being part of a given functional class or not. To be conservative, we set strict thresholds for identifying predictive grid cells, particularly in the path-integrating RNNs. This may contribute to why the number of predictive grid units was *≈* 5%. Recently, there has been a growing consensus in the spatial neuroscience community that such classification is overly limiting and subjective. Instead, measuring properties of spatially-tuned cells (e.g., grid score, spatial information) enables a more holistic understanding spatial representations. Currently, there does not exist a single scalar metric that measures the strength of predictive grid tuning. We attempted to provide one based on the relative grid score improvement, but we recognize that this itself is somewhat limited (e.g., a cell can have a relative grid score improvement of 1.0 by having its grid score increase from 0.01 to 0.02). Creating metrics that more completely characterize predictive grid cells will help resolve some of these challenges.

Finally, similar predictive grid tuning in MEC and RNNs do not necessarily establish identical computational roles. Indeed, while we find a similar number of retrospective grid units as predictive grid units in all 10 independently trained RNNs (Fig. 3), analysis of MEC data finds much less evidence of retrospective grid coding [26] (Fig. S4). This points to important differences between MEC and RNNs. In addition, the architecture we used for the path-integrating RNNs lacks aspects of biological realism, in contrast to other attempts to model predictive grid cells [44, 45]. Despite this, the analysis of path-integrating RNNs makes predictions about predictive grid cell function, which was tested on MEC sleep data and can be further probed with future experiments.

### What is the computational role of predictive grid cells?

If predictive grid cells are not used directly for planning, what are they useful for? One possibility is that, being part of the grid cell circuit, they play a role in shaping the grid cell dynamics, an idea consistent with recent findings showing that a predictive signal is needed to maintain attractor dynamics in the head-direction network of head-fixed mice [49]. To test this, we performed a preliminary analysis where we ablated predictive grid units in path-integrating RNNs and examined the effect their removal had on the population activity of grid units (see Methods, for more details). We found a subtle affect, with networks that had predictive grid units ablated having intact grid unit autocorrelation at short time-scales, but increased autocorrelation at longer time-scales (Fig. S11B). This was unique to predictive grid units, as removing other functional classes did not lead to the same effect (Fig. S11C–E). We hypothesize that predictive grid units are used to “repel” the grid unit activity away from where it currently is along the attractor. This would be useful when moving through space, since – until one has traveled half the length of a grid spacing – the grid unit activity should become increasingly distinct.

### Future work

While we have showed that predictive grid coding can emerge without explicit planning, it remains to be tested whether predictive grid coding becomes stronger and/or more prevalent when path-integrating RNNs are used for goal-based navigation [6]. Similarly, whether predictive grid units can be leveraged for improved planning is an important question future work should address. Indeed, while left-right-alternating theta-sweeps in MEC have been argued to be part of the local circuit [24], they have also been found to be modulated by the specific features of the environment and by demands of a given task [25] that may be relevant for planning. This suggests that regions downstream of MEC and hippocampus could still leverage this signal for planning, which may also hold true for predictive grid cells. To better understand how predictive grid cells might shape grid unit activity, particularly through continuous attractor network dynamics, mechanistic models that include predictive grid cells can be developed. By changing the path statistics the RNNs are trained on, it may be possible to identify what aspects of path-integration most strongly drive the emergence of predictive grid cells [50]. New neurophysiological experiments can be performed to examine whether predictive grid coding is modulated by manipulations that are known to affect grid coding, such as HCN1 channel knockout [51]. Finally, predictive grid cells should be looked for in electrophysiological recordings of other animals that exhibit grid cells, such as bats [52, 53].

## Methods

### Electrophysiology recordings

The MEC data analyzed in this paper comes from a publicly available data set, which has previously been described in detail [27]. We provide brief summary of the methodology and the experimental paradigms used during the recordings.

Three male rats (Long Evans – Rats Q, R, and S) were implanted with Neuropixels silicon probes [37, 54]. These probes were targeted at the MEC-parasubiculum region and the surgery was performed as described previously [35, 37]. After three hours of recovery, recordings were performed. In the recordings analyzed, the rats foraged for randomly dispersed corn puffs in a 1.5 *×* 1.5 m^2^ square open field arena, with walls of height 50 cm. The rats were familiar with the environment and task, having trained 10–20 times prior to the implantation. The rats were food restricted to motivate their foraging, being kept at a minimum of 90% of their original body weight (300–500 grams).

All procedures in the original study were approved by the Norwegian Food and Safety Authority and done in accordance with the Norwegian Animal Welfare Act and the European Convention for the Protection of Vertebrate Animals used for Experimental and Other Scientific Purposes. Protocols were approved by the Norwegian Food Safety Authority (FOTS ID 18011 and 18013).

### Electrophysiology post-processing

The MEC data analyzed in this paper was post-processed, before made publicly available. We describe, in brief, the post-processing performed [27], as well as the post-processing we performed on the downloaded data. Spike sorting, via KiloSort 2.5 [37], was applied to the data recorded from the Neuropixel probes. Individual units were deemed putative cells if their average spike rate was in the range of 0.5–10Hz, and 99% of their interspike intervals were greater than 2 ms.

For each putative cell, ratemaps were constructed by averaging the activity at binned spatial positions in the open field arena. This raw ratemap was smoothed, using a Gaussian kernel. The autocorrelation of these ratemaps were computed, and a grid score calculated, as described previously [55]. Grid module identities were assigned by first projecting the 2D autocorrelogram of every recorded unit onto a 2D space using the non-linear dimensionality reduction algorithm UMAP [56]. Then, the DBSCAN algorithm [57] was used to cluster the units, based on their position in the dimensionally reduced space. Cluster membership served as the basis for grid module classification, with the largest cluster being removed as it was found to not contain spatially selective autocorrelograms. This non-supervised module assignment yielded clusters of high grid scores and similar grid spacing and orientation within each cluster-module. More details can be found in Gardner et al. (2022) [27]. Only units that belonged to an identified grid module were included in the dataset. We performed no further analysis regarding module identity.

From the downloaded spike trains, we constructed ratemaps and spatial autocorrelograms (SACs) in a similar manner, using code made publicly available [6, 13].

### MEC temporal-shift analysis

To identify predictive grid cells from the MEC data using time-based prospective shifts, neural activity was aligned with the animal’s position across a range of temporal offsets, Δ. For each cell and temporal offset, a shifted firing-ratemap was calculated as

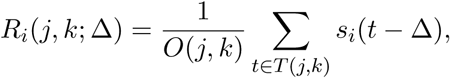

where *s_i_*(*t*) denotes the spiking activity of cell *i* at time *t*, *T* (*j, k*), denotes the time points at which the animal occupied spatial bin (*j, k*), and *O*(*j, k*) is the corresponding occupancy. At Δ = 0, this is equivalent to the unshifted firing-ratemap, as *s_i_*(*t −* Δ) = *s_i_*(*t*). SACs were computed from each temporally shifted firing-ratemap, and grid score was calculated using the same six-fold symmetry measure referenced above [55], producing a grid-score curve *GS*(Δ) for each cell.

Temporal offsets were evaluated in both prospective and retrospective directions. Under the convention used here, positive Δ values correspond to prospective shifts and negative Δ values correspond to retrospective shifts. For each cell, the temporal offset producing the maximum grid score, Δ_max_, was identified from resulting *GS*(Δ) curve.

### Predictive grid cell classification

An increase in grid score following temporal shifting does not, by itself, establish genuine predictive coding, because apparent improvements may arise from variability in neural firing, behavioral sampling, or chance alignment between spike trains and position. To distinguish genuine temporal predictive coding from such effects, we performed a circular-shift shuffle analysis.

For each recording, 100 surrogate datasets were generated by applying randomly selected circular shift to the animal’s x- and y-position trajectories, while leaving the recorded spike trains unchanged. This procedure disrupted the temporal alignment between neural activity and position while preserving the overall trajectory and sampling structure of each recording. The same grid-score analysis used for the original data was then applied to each shuffled realization.

For each cell, the maximum grid score in the original recording, *g* = max_Δ_ *GS*(Δ), was compared with the grid score obtained under each shuffled realization, *g̃_i_* = *GS*_shuffle_(0) for *i* = 1*, …,* 100. The number of shuffled realizations whose maximum grid score exceeded the observed maximum was defined as

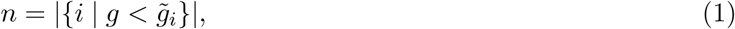

where *| · |* denotes the cardinality of the set. A cell was considered statistically significant under the shuffle analysis when fewer than 5 of the 100 shuffled realizations produced a grid score greater than the experimentally observed value, *n <* 5.

Statistical validation was combined with three additional criteria for a cell to be classified as a predictive grid cell. In particular, a cell must:

- Exhibit weak grid structure when no shift is present – *GS*(0) *<* 0.5
- Achieve its maximum grid score at a prospective temporal offset – Δ_max_ *>* 0
- Exhibit a sufficiently strong maximum grid score – *GS*(Δ_max_) *>* 0.3.

Thus, predictive grid cells were defined as neurons satisfying all three of the above criteria and the shuffle-based statistical criterion simultaneously. This *GS*(0) threshold excluded neurons that were already strongly grid-like at the animal’s current position, while the *GS*(Δ_max_) threshold required the predictive effect to reach a minimum level of grid-like organization. The shuffle criterion further required that the observed increase in grid score be unlikely to result from chance temporal alignment.

### MEC spatial-shift analysis

To test whether predictive grid structure could also be identified through spatial (instead of temporal) projection, the animal’s recorded position trajectory was shifted forward and backward along its instantaneous movement direction, while neural spike timing was kept unchanged. At each time point, the direction of movement was calculated from the x*−* and y*−*coordinates, and the position was displaced by a signed distance *D*:

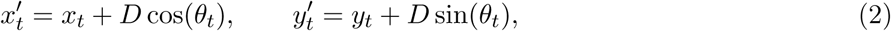

where *D* represents the spatial projection distance and *θ* is the animal’s instantaneous movement direction, 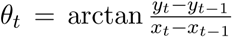 . Positive values of *D* projected the animal’s position forward along its trajectory, whereas negative values projected it backward. Spatial projection distances ranged from *−*0.50 to +0.50 m in 0.02 m increments.

For each spatial projection, positions that fell outside the original 1.5 *×* 1.5 m^2^ recording arena were excluded, together with the corresponding time points from each cell’s aligned spike train. Grid scores were then recomputed using the same spatial firing-ratemap and SAC analysis described above, producing a grid-score curve *GS*(*D*) for each cell. Cells were considered predictive grid cells (via a spatial-shift) when their maximum grid score occurred at a positive spatial projection distance and exceeded the minimum grid-score threshold used for predictive-cell identification. This complementary analysis revealed spatially predictive grid structure across the recording modules (Figs. S2, S3).

### Path-integrating recurrent neural networks

We trained 10 independently initialized recurrent neural networks (RNNs) to perform pathintegration in a 2.2 *×* 2.2 m^2^ square arena [6–8]. Each network represented position with 512 difference-of-Gaussian place cells (center width, 0.12 m; surround ratio, 2) and contained 4096 recurrent rectified linear units. A learned, bias-free encoder initialized the hidden state from the place-cell code at the starting position. The network then updated this state from two-dimensional velocity input, and a bias-free linear decoder predicted the current place-cell distribution. We optimized cross-entropy loss with an *ℓ*_2_ penalty on the recurrent weights (*λ* = 10*^−^*^4^) using Adam (learning rate, 10*^−^*^4^) with 200 trajectories per batch and 1000 batches per epoch, for 100 epochs. Our networks shared hyperparameters but often differed in their initial weights and sampled trajectories.

Training trajectories were 800-ms pseudo-random walks. Speed and heading changed smoothly, and agents slowed and turned inward near arena boundaries. The network received its initial position and instantaneous velocity but no goal, planned route, or future-position signal. We also evaluated the trained networks, without retraining, on trajectories with fixed direction and speed, and the same spatial-tuning and classification procedures were applied to both trajectory types.

Importantly, our RNNs were consistent in all choices of architecture and hyper-parameters as described previously by Sorscher et al. (2022) [8]. The sole exception being we trained on trajectories that were twice as long as the ones they trained on (800 ms compared to the 400 ms used by Sorscher et al.).

### Predictive grid unit classification

To identify predictive grid units in the RNN, we performed a classification approach similar to the one employed on the MEC data. In particular, we aligned the RNN agent’s position to the recurrent units’ activity at different temporal offsets, computing the grid score from the resulting ratemap and SAC across a range of temporal-shifts (Δ from *−*400 ms to 400 ms). Shifts were applied within trajectories only.

We first began by identifiy all units that could be classified as a standard grid units. This classification was done by finding units whose *GS*(0) *>* 0.3 and whose *GS*(0) was statistically significant, compared to the shuffle analysis (as described previously). In order for a unit to be classified as a predictive grid unit, it must:

Exhibit weak grid structure when no shift is present – it must not be classified as a standard grid unit
Achieve its maximum grid at a prospective temporal offset – Δ_max_ *>* 0
Exhibit a sufficient strong maximum grid score – *GS*(Δ_max_) *>* 0.3 and *GS*(Δ_max_) must be statistically significant compared to the shuffle analysis
Have smoothly varying *GS*(Δ) curves – *GS*(Δ_max_) must not be more than 1.5*×* greater than the next largest value of *GS*(Δ).

Retrospective units were identified using the same criteria, with Δ_max_ *<* 0 instead.

In order to assess path-dependence, within each network we identified pairs of trajectories that occupied the same 0.05 *×* 0.05 m^2^ spatial bin at the same time step and computed population-vector correlations separately for the standard and predictive grid-unit populations. We related these correlations to the Euclidean difference between the paired velocity vectors, ||**v**_1_ – **v**_2_||_2_, and then summarized them within velocity-difference bins for visualization.

### Band and border analyses

We computed border and band scores using methods previously developed [1, 32], which we briefly summarize here. Band score was the maximum correlation between a unit’s ratemap and two-dimensional cosine templates with spatial frequencies from 0 to 2 cycles/m in increments of 0.1 cycles/m [32]. Even though we excluded the “constant” template, we kept the templates with one zero-frequency dimension. Border score was defined as (*C_M_ − D_M_* )*/*(*C_M_* + *D_M_* ) [1], where *C_M_* is the maximum wall coverage of a firing field and *D_M_* is its firing-rate-weighted distance from the nearest wall. Firing fields were connected regions with activity above 30% of peak. Within each network, units above the 95^th^ percentile of the relevant score distribution were classified as band or border units. We measured their overlap with standard, predictive, and retrospective grid populations and compared their maximal prospective and retrospective grid-score increases.

### Toroidal ratemaps

To construct the toroidal ratemaps used in Fig. 6, we use open source code that was published alongside Gardner et al. (2022) [27], which was accessed at https://github.com/erikher/GridCellTorus. Briefly, to construct the toroidal ratemaps, topological data analysis (TDA) was performed on the neural activity. Importantly, any cell that was found to be a conjunctive head-direction grid cell was excluded from all subsequent analysis. We used the same identification as Gardner et al., which was available on the Github repository. We found little overlap between predictive grid and conjunctive cells.

As described in Gardner et al., the neural activity was first dimensionally-reduced via PCA, with 6 components being kept. On this lower dimensional subspace, persistent cohomology was used to generate a barcode, which describes the emergence and disappearance of topological structures (e.g., circles, two-dimensional cavities). From this barcode, a complex process was used to find a parametrization of the underlying torus, *ϕ*_1_*, ϕ*_2_ *∈* [0, 2*π*). We refer the interested reader to the original paper [27]. Having found *ϕ*_1_ and *ϕ*_2_ for all time-steps, we then binned the toroidal angles and computed the average activity of each MEC cell. We find qualitatively similar ratemaps as was reported in Gardner et al. To quantify the amount of selectivity predictive grid cells encoded about *ϕ*_1_ and *ϕ*_2_, we computed the spatial information (SI) [36],

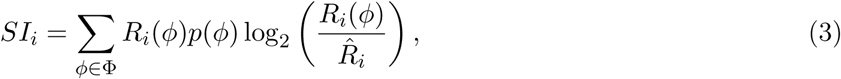

where Φ is the binned toroidal angles, *R_i_*(*ϕ*) is the average firing rate of cell *i* at binned toroidal angle *ϕ*, R^i is the average firing rate of cell *i*, and *p*(*ϕ*) is the probability of the toroidal angle being *ϕ*.

We constructed shifted toroidal ratemaps using the same process as described earlier, shifting the activity Δ ms with respect to the toroidal angle, and then re-computing the ratemaps and the SI.

### Ablation analysis

We separately ablated predictive grid, retrospective grid, and band units by setting all incoming and outgoing weights to 0. This has the effect of fixing their activations to 0 at every time step. We also ablate a random subset of “non-dead” units (i.e., units that exhibit activity in at least 10% of the spatial bins of their ratemaps). The total number of units ablated is matched to be the same as the number of predictive grid units we ablate. We perform 20 random ablations and take their average effect on the autocorrelation. In all cases, we ensure that we do not ablate standard grid units, as this is the population we are computing the autocorrelation on. For predictive and retrospective grid units, this is ensured by the fact that there is no overlap between the two populations. For band units, we remove any unit that is in the intersection between the units classified as band units and those classified as standard grid units. We then ablate the remaining. Finally, for the random ablations, we randomly sample from units that are both non-dead and not classified as grid units.

We performed PCA on grid unit activity from 100 trajectories from the original, intact network, and then utilized the first two components. Base and ablated activity was projected into this fixed basis. PCA was not refitted after we performed the ablation (Fig. S11A). Therefore, any changes in the projected trajectories reflected changes relative to the intact population geometry, instead of just rotations between separately fitted spaces.

To compute the autocorrelation, we used the cosine similarity, which is a normalized measure of similarity. This analysis tested whether ablations caused population trajectories to return toward previously visited states, rather than simply reducing their activation rate (Fig. S11B–E).

## Acknowledgments

We thank Kevin Sit, Nora Wolcott, Santiago Acosta-Mendoza, Mikkel Lepperød, and Chris Cueva for helpful discussion. We thank Santiago Acosta-Mendoza for the code, in part, used to analyze the MEC data (published previously [13]). W.T.R. is supported by the Data Science and AI Institute (DSAI) at Johns Hopkins University.

## Data and materials availability

All code used for the analysis presented in this paper is publicly available: https://github.com/Dynamical

-Intelligence-Group/predictive-grid-cells. The MEC recordings analyzed was accessed from the publicly available repository: https://figshare.com/articles/dataset/Toroidal_topology_of_population_activity_in_grid_cells/16764508?file=35078602.

## Author contributions

WTR conceived of the study. OCF and WTR analyzed the MEC data. AS and WTR analyzed the RNNs. AS and WTR decided on and implemented analysis. WTR made all figures. AS and WTR wrote original draft of paper. All authors contributed to final draft.

## AI usage statement

Generative AI was used for assisting in some (but not all) code generation, debugging, and initial writing. In addition, generative AI was used for explaining unfamiliar concepts/code. However, all finished code, writing, and figure making was fully validated by the authors and we take responsibility for all statements made.

^1^Using a looser criteria for classifying predictive grid units, we find closer to 10% of being classified, with similar tuning profiles as found in Fig. 2.

^2^One of the nine recordings in the dataset had only a single predictive grid cell. Therefore, we exclude it from our analysis

^3^No sleep data was collected on day 1 of recording rat R, ruling out R11 and R12.

---

**Figure S1:**
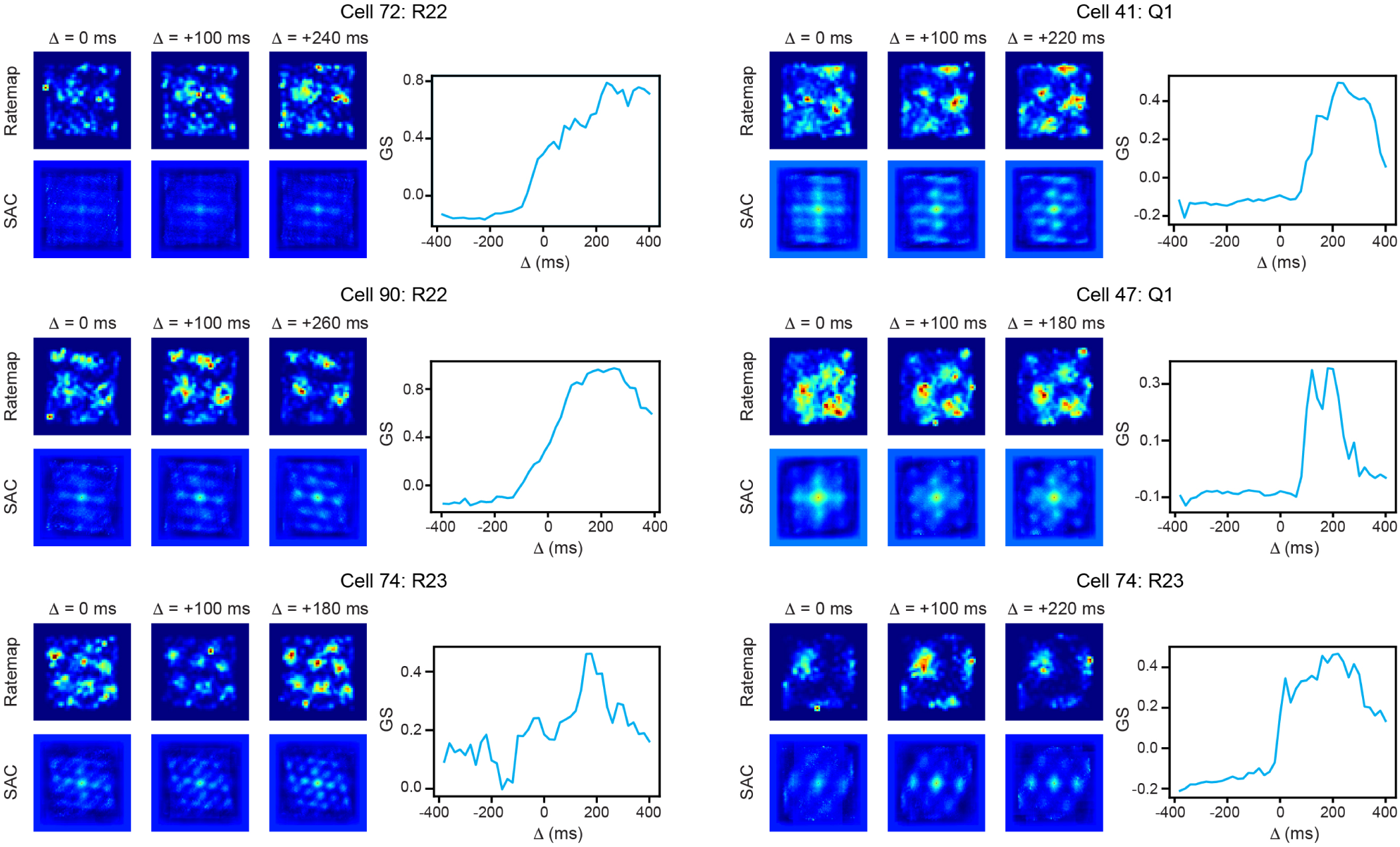
Additional examples of cells classified as predictive grid cells from Gardner et al. (2022) dataset.

**Figure S2:**
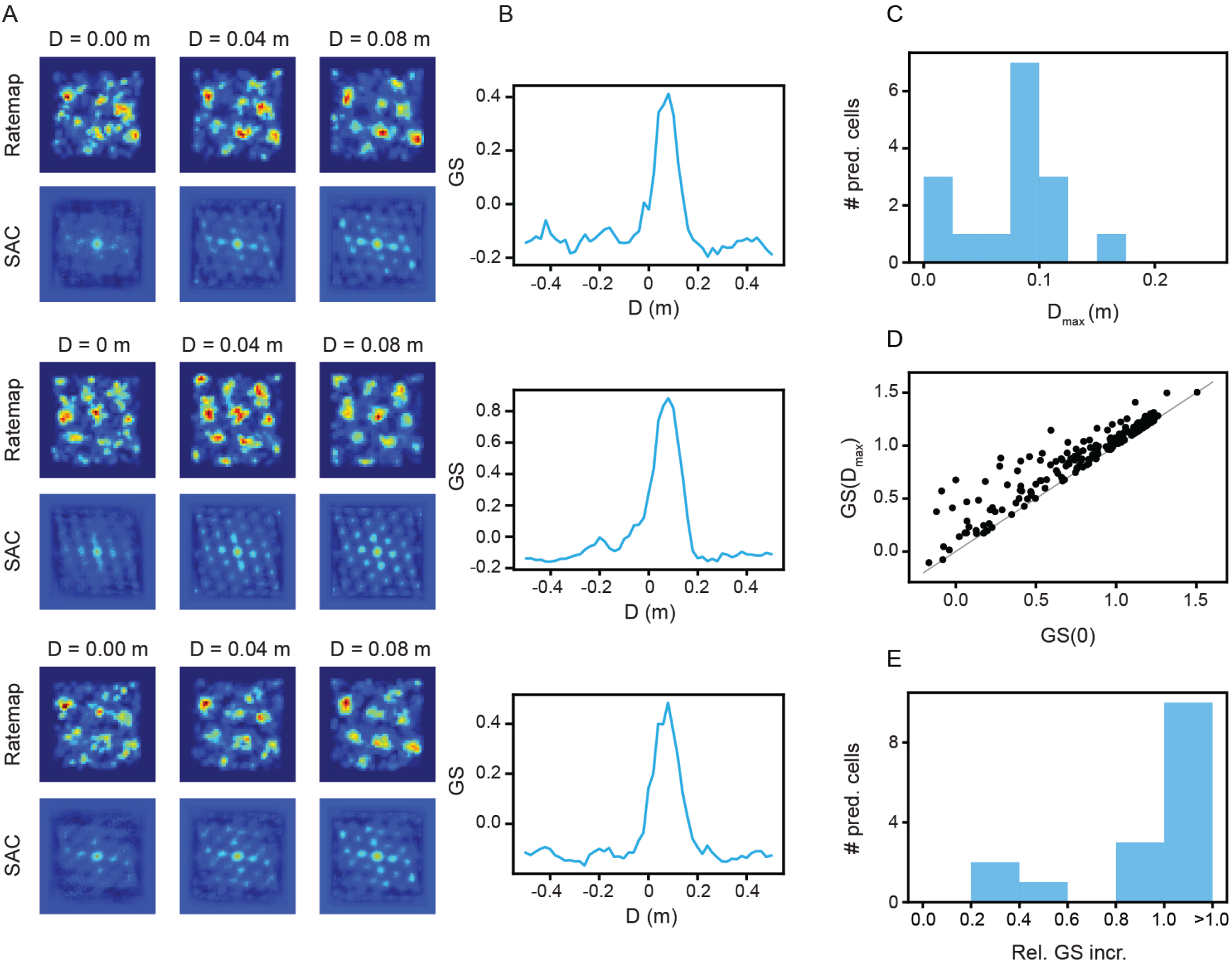
Predictive grid cells, identified with spatial shifting, are present in Gardner et al. (2022) MEC recording (recording ID R21). (A) Example ratemaps and spatial autocorrelograms (SACs), at different spatial shifts, for cells classified as predictive grid cells. (B) Grid score as a function of spatial shift tuning curves for all three predictive grid cells in (A). (C) Distribution of spatial shifts with maximal grid score (Δ_max_) for all cells classified as predictive grid cells. (D) Grid score at *D* = 0 m, compared to grid score at *D* = *D*_max_, for all recorded cells. grid score at *D* = 0 m divided by grid score at Δ = *D*_max_ for all recorded cells. (E) Distribution of relative grid score increase for all cells classified as predictive grid cells.

**Figure S3:**
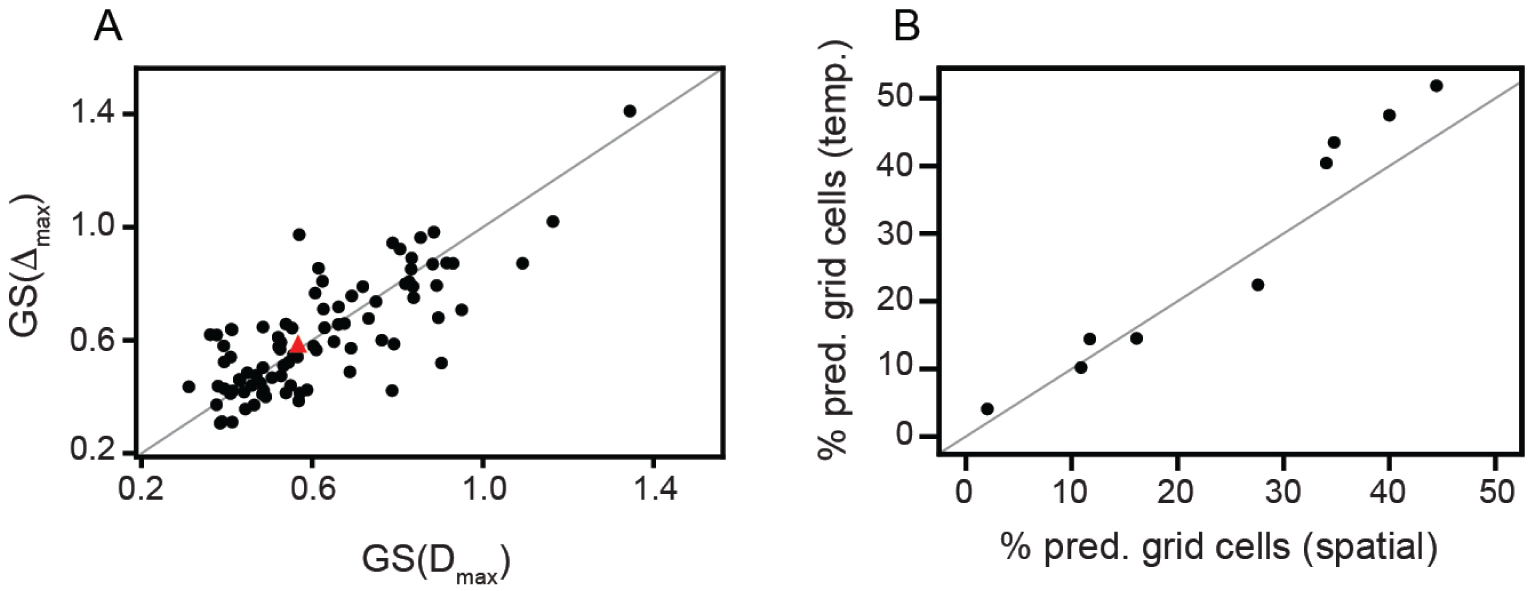
Comparison between of predictive grid cells found using temporal and spatial shifting. (A) Maximum grid score of all cells identified as predictive grid cells, when shifting in space (x*−*axis) versus shifting in time (y*−*axis). Red triangle is median value across the population. (B) Percent of non-grid MEC cells that were classified as predictive grid cells when shifting in space (x*−*axis) versus shifting in time (y*−*axis). Each dot represents one recorded grid module. (A)–(B) Gray line corresponds to *x* = *y*.

**Figure S4:**
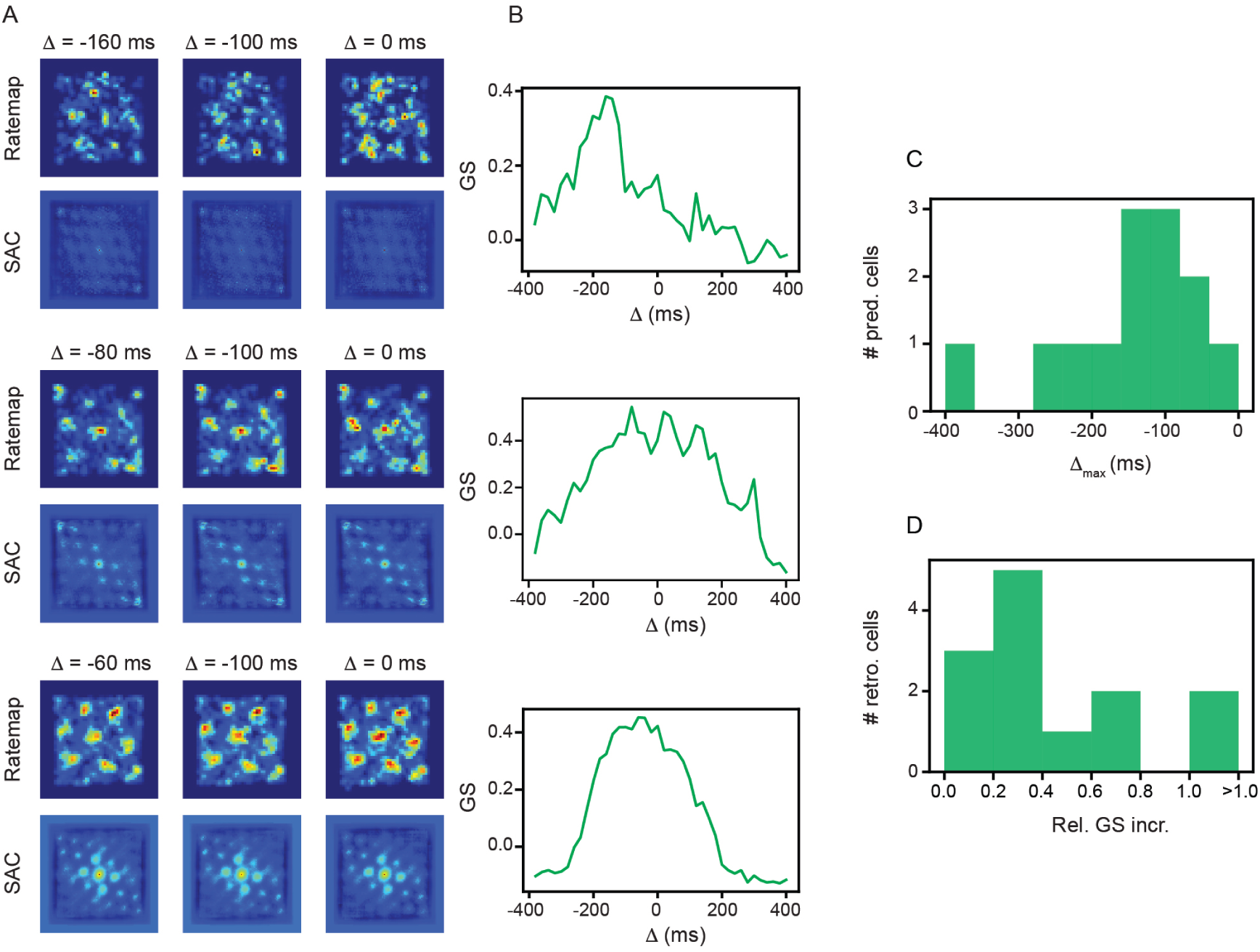
Weak retrospective grid coding exists in Gardner et al. (2022) MEC recording (recording ID R21). (A) Example ratemaps and spatial autocorrelograms (SACs), at different temporal shifts, for cells classified as retrospective grid cells. (B) Grid score as a function of temporal shift tuning curves for all three retrospective grid cells in (A). (C) Distribution of temporal shifts with maximal grid score (Δ_max_) for all cells classified as retrospective grid cells. (D) Distribution of relative grid score increase for all cells classified as retrospective grid cells.

**Figure S5:**
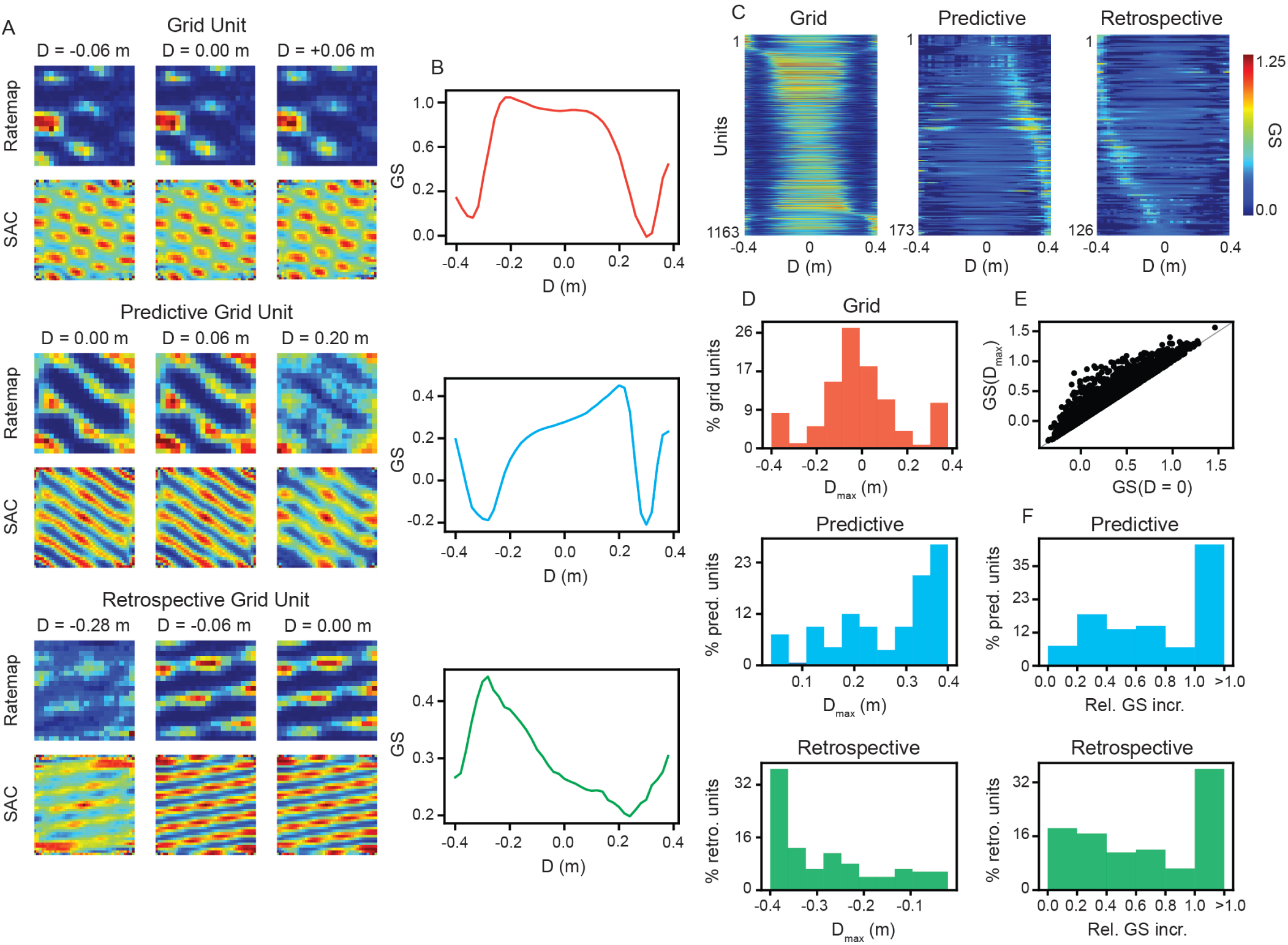
Predictive and retrospective grid units, identified with spatial shifts, are present in RNNs trained to perform path-integration. Same as Fig. 2, but when using a spatial shift.

**Figure S6:**
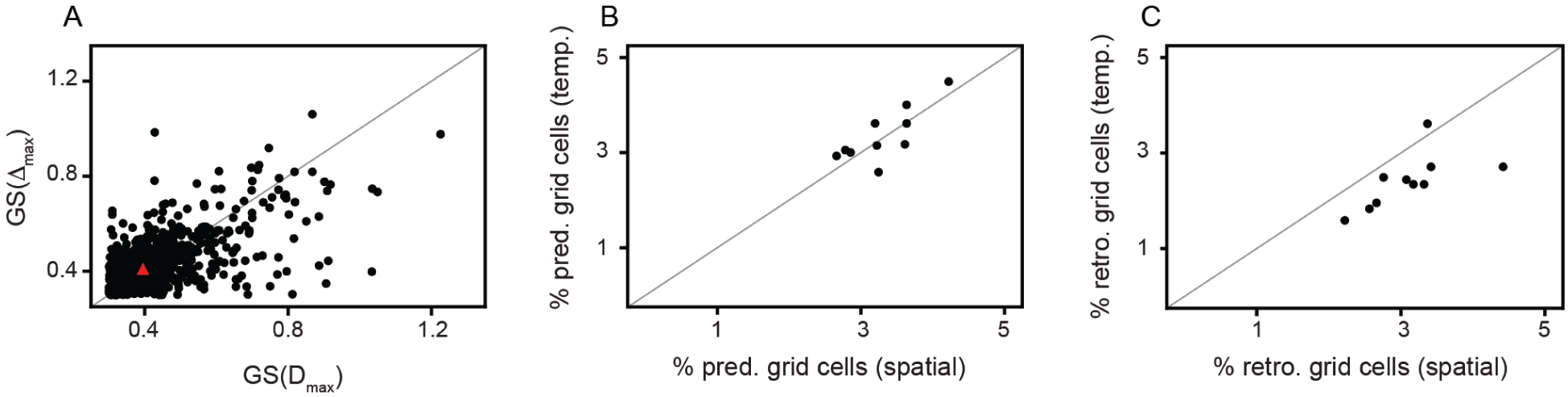
Comparison between of predictive grid units found using temporal and spatial shifting. (A) Maximum grid score of all cells identified as predictive grid units, when shifting in space (x*−*axis) versus shifting in time (y*−*axis). Red triangle is median value across the population. (B) Percent of non-dead units that were classified as predictive grid units when shifting in space (x*−*axis) versus shifting in time (y*−*axis). Each dot represents one independently trained RNN. (A)–(B) Gray line corresponds to *x* = *y*.

**Figure S7:**
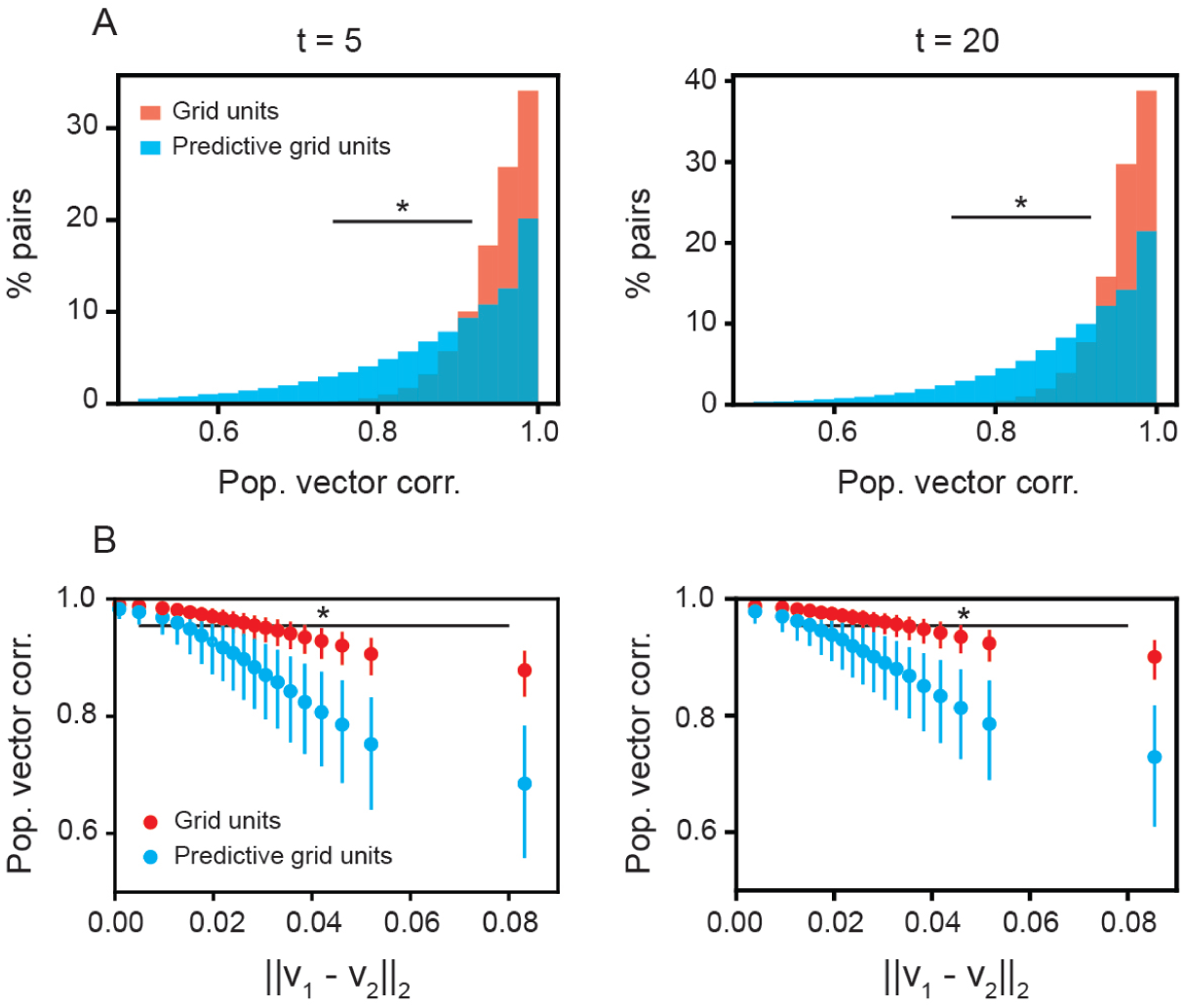
Predictive grid units are more future path-dependent than grid units, across different values of *t*. (A) Same as Fig. 4B, but for different values of *t*. (B) Same as Fig. 4D, but for different values of *t*.

**Figure S8:**
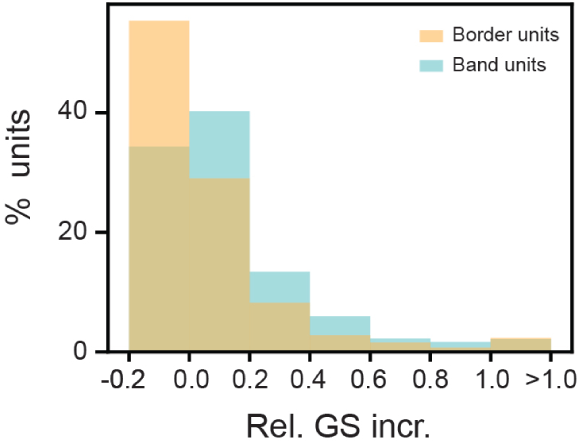
Band units exhibit greater relative grid score increase for prospective shifts than border units. Same as Fig. 5G, but for border and band units.

**Figure S9:**
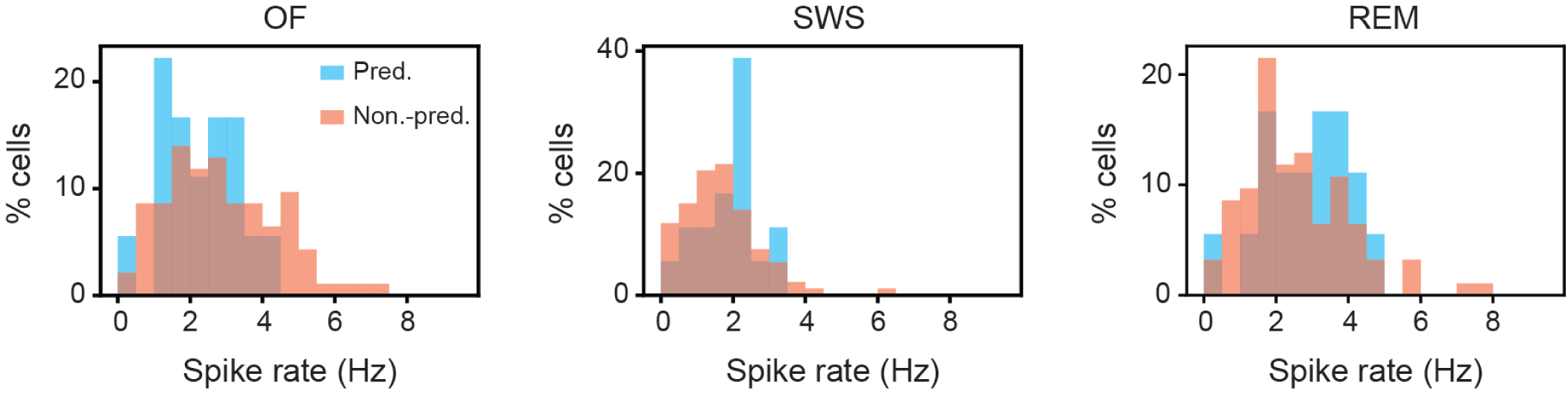
Predictive grid cells do not have statistically significantly distinct firing distribution from non-predictive grid cells, including during sleep.

**Figure S10:**
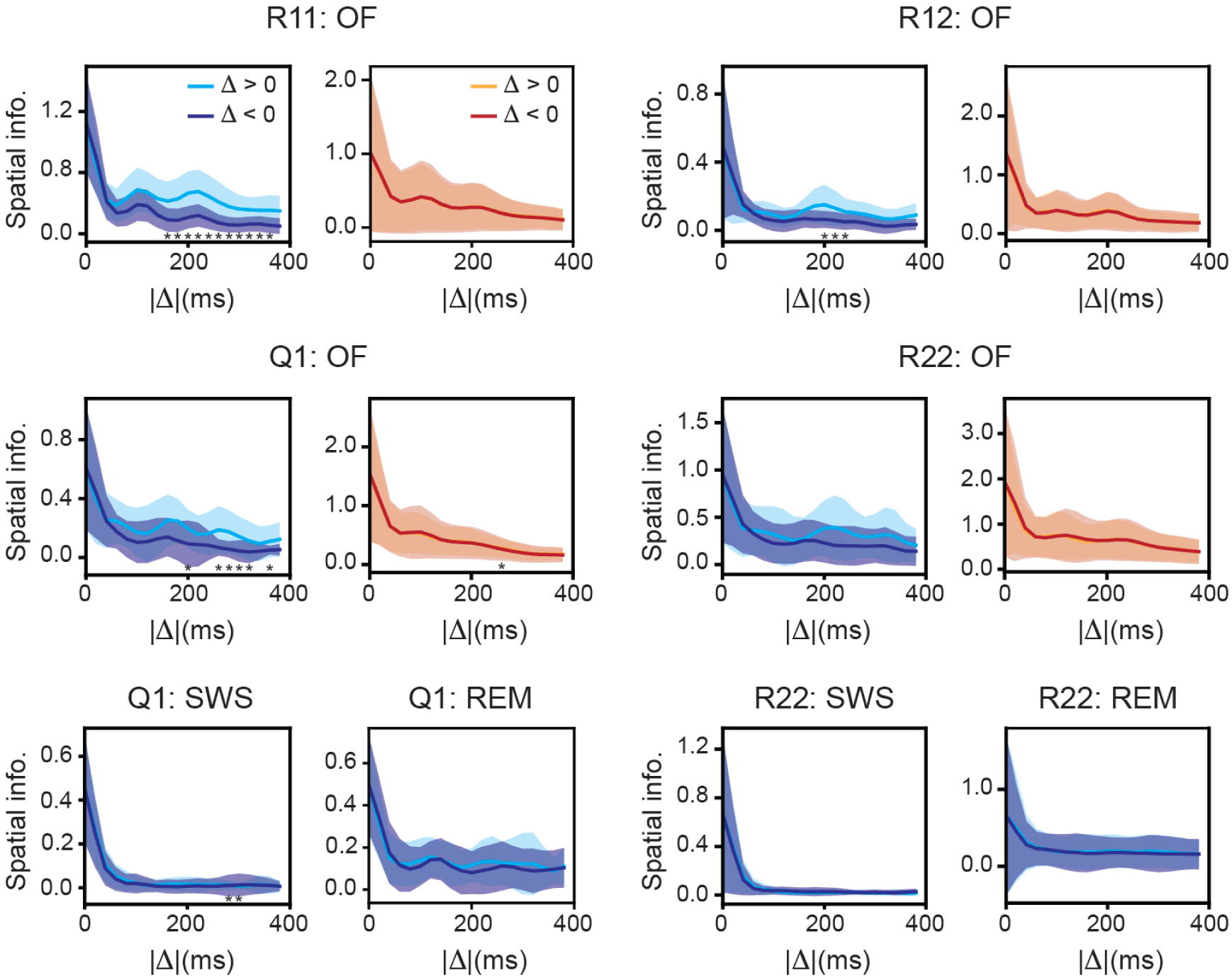
Additional toroidal ratemap spatial information vs. temporal shift, for recordings from Gardner et al. (2022) dataset.

**Figure S11:**
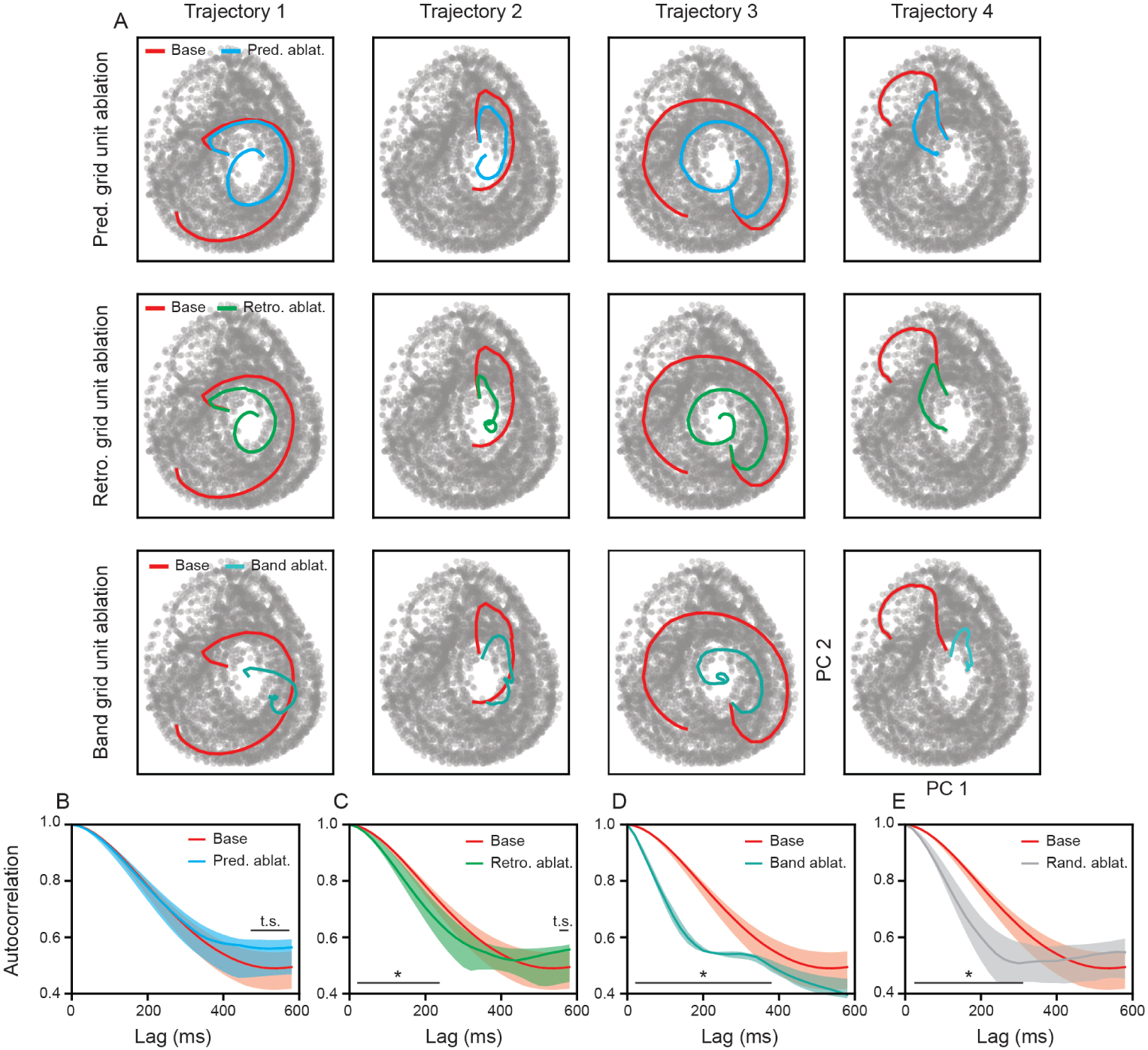
Predictive grid units shape grid cell population activity by long time-scale de-correlation. (A) Example standard grid unit population activity trajectories, projected along the top two principal components (PCs), with and without predictive grid, retrospective grid, and band units ablated. Red line denotes trajectory of network. Blue, green, and cyan lines denote trajectory with ablation. Gray dots denote the population activity from base network used to compute the PCs. (B)–(E) Autocorrelation, as a function of temporal lag, of standard grid unit population with and without predictive grid, retrospective grid, band, and random units ablated, respectively. Solid line denotes median and shaded area denotes 25^th^ to 75^th^ percentile. Star denotes *p*-value *<* 0.05 using one-sided KS test and t.s. denotes *<* 0.05 *p*-value *<* 0.10 using one-sided KS test.

